# The first chicken oocyte nucleus whole transcriptomic profile defines the spectrum of maternal mRNA and non-coding RNA genes transcribed by the lampbrush chromosomes

**DOI:** 10.1101/2024.02.05.577752

**Authors:** A. Krasikova, T. Kulikova, M. Schelkunov, N. Makarova, A. Fedotova, V. Berngardt, A. Maslova, A. Fedorov

## Abstract

Lampbrush chromosomes, with their unusually high rate of nascent RNA synthesis, provide a valuable model for studying the mechanisms of global transcriptome up-regulation. Here, we aimed to establish a full set of sequences transcribed on the lateral loops of chicken lampbrush chromosomes. For the first time, a whole-genomic profile of transcription along the entire length of all lampbrush chromosomes in the chicken karyotype including sex chromosomes and dot chromosomes was obtained. For that, we performed RNA-seq of oocyte nuclear and cytoplasmic total RNA, poly(A) RNA and small RNA libraries and aligned expressed RNA sequences against the chicken genome. This led to the identification of a full spectrum of maternal RNAs that accumulate in the nucleus and cytoplasm of chicken oocytes at the lampbrush chromosome stage, are then transferred to the zygote and can be used during the early stages of embryogenesis, including transcripts of protein-coding genes, long non-coding RNAs and small housekeeping non-coding RNAs. We also present the first high-throughput transcriptome characterisation of miRNAs and piRNAs in chicken oocytes at the lampbrush chromosome stage. Major targets of predicted piRNAs include CR1 and LTR containing retrotransposable elements. Transcription of tandem repeat arrays including 41bp higher-order repeats as well as unique centromere sequences was demonstrated by alignment against the whole telomere-to-telomere chromosome assemblies. We show that transcription of telomere-derived RNAs, including telomeric repeat-containing RNA (TERRA) and subtelomeric repeat-containing RNA, is initiated at adjacent LTR elements. With nuclear RNA-seq, we obtained information about a wider set of transcripts, including long non-coding RNAs retained in the nucleus and stable intronic sequence RNAs (sisRNAs). We identified 5’ UTR sisRNAs that may potentially support host gene transcription both during oogenesis and after activation of the embryonic genome. For a number of protein-coding genes, we visualised nascent transcripts and demonstrated their co-transcriptional splicing on the lateral loops of lampbrush chromosomes by RNA-FISH. The nuclear RNA-seq profile predicted the chromomere-loop organisation of the genomic regions. During oocyte maturation, transcriptional unit boundaries were maintained, while transcriptional output tended to decrease. We conclude that cytoplasmic and nuclear transcripts emerge from nascent transcripts on the lateral loops of lampbrush chromosomes. Most of gene transcripts are initiated at promoters, normally spliced, terminated and polyadenylated. The set of genes transcribed on the lampbrush chromosomes is required for basic cellular processes, is characterised by a broad expression pattern and is similar to sets of genes expressed in other hypertranscriptional systems. We conclude that hypertranscription on the lateral loops of giant lampbrush chromosomes is the main mechanism for synthesising large amounts of transferred to the embryo maternal RNA for thousands of genes.

## 1. Introduction

Factors accumulated during gametogenesis, specifically oogenesis, play a key role in regulating the embryonic genome. Maternal RNAs produced and stored in the oocyte serve both as RNA templates for protein synthesis and non-coding regulators of embryonic gene expression. Maternal transcripts are required for oocyte maturation, fertilisation and early development (Tora and Vincent 2021). The oocyte cytoplasmic protein-coding RNAs can either be actively translated or be stored as dormant maternal transcripts for use after fertilisation. The transcriptomes of the whole oocytes are being studied intensively. However, the source of maternal RNAs accumulated in the oocyte cytoplasm is difficult to identify during these studies. In addition, due to the high concentration of mature mRNA for a number of protein-coding genes, less abundant regulatory RNAs may escape the focus of investigation.

All vertebrates, except placental and marsupial mammals, have a hypertranscriptional type of oogenesis, accompanied by an extremely high rate of RNA synthesis in the oocyte nucleus (also called germinal vesicle) (Gall 2014). The high stability and constant renewal of the maternal RNA pool creates the conditions for the long-term maintenance of egg viability in the ovary. During the hypertranscriptional stage of oogenesis, in fishes, reptiles, birds, some insects and probably monotremes, the chromosomes acquire a lampbrush morphology (Callan 1986; Macgregor 2013; Gall 2014). Lampbrush chromosome chromatin is distinctly structured, with a predetermined chromomere-loop configuration. Active transcription of various genomic sequences takes place on the lateral loops, also called transcription loops (Morgan 2018; Mirny and Solovei 2021; Krasikova *et al*. 2023a). The extraordinary length of the lateral loops and the density of newly synthesised RNA make it possible to visualise and map individual transcription units with exceptional resolution (Kaufmann *et al*. 2012; Kulikova and Krasikova 2022).

The study of transcripts found in oocytes with the hypertranscriptional type of oogenesis began in the sixties of the last century. It was shown that about 3% of the genome is transcribed on the *Xenopus laevis* lampbrush chromosomes. Nonribosomal RNA synthesised during the lampbrush chromosome stage is retained throughout oogenesis; these maternal template-active RNAs “must ultimately be utilised in embryogenesis” (Davidson *et al*. 1966). Hybridisation experiments of newly synthesised total RNA from *Xenopus* lampbrush stage oocytes with non-repetitive genomic DNA revealed high complexity and “enormous informational diversity” of the oocyte transcriptome (Davidson and Hough 1969). Similar hybridisation experiments with complementary DNA derived from oocyte poly-A mRNA demonstrated that pre-mRNA sequences are present in both *Xenopus* and *Triturus* oocyte nuclear RNA fraction and estimated that about 3% of nuclear RNA becomes mRNA, giving rise to about 10000 different mRNA species (Davidson and Hough 1971; Sommerville and Malcolm 1976).

In 2012, a group led by John Gurdon characterised the transcriptome of *Xenopus tropicalis* lampbrush stage oocytes using high-throughput sequencing (Simeoni *et al*. 2012). The authors described transcript content for over half of the 20000 annotated genes, including hundreds of correctly spliced transcripts of cell type-specific genes that are usually expressed during development.

Given the differences in RNA profiles between the nucleus and cytoplasm, it has been suggested that RNA sequencing of separate nuclear and cytoplasmic RNA fractions provides a more accurate picture of gene expression and RNA processing (Zaghlool *et al*. 2013). Nuclear retention of transcripts can be used to prevent them from being translated (Butto *et al*. 2023). Moreover, nuclear RNA fractions are also enriched in long non-coding RNAs and partially spliced nascent transcripts, reflecting primary transcript levels (Mitchell *et al*. 2012). After oocyte nuclear envelope breakdown, nuclear transcripts are also released into the cytoplasm and thus can be transferred to the zygote (Davidson *et al*. 1966). Joseph G Gall and co-authors were the first to sequence the complete transcriptomes of isolated nuclei and cytoplasm of *X. tropicalis* oocytes (Gardner *et al*. 2012). In this work, transcripts of about 6700 protein-coding genes, mainly represented by exon sequences, were identified in the oocyte cytoplasm. At the same time, the transcriptome of the oocyte nuclei was enriched for the intron sequences of the same genes. A number of intron-containing transcripts showed unusually high stability and persisted until the mid-blastula stage of embryonic development, when the *Xenopus* zygotic genome is activated (Gardner *et al*. 2012). However, other types of regulatory RNAs, including long non-coding RNAs and short regulatory RNAs in the *Xenopus* oocyte nucleus, have not been characterised.

In the chicken (*Gallus gallus domesticus*), high transcriptional output also leads to transformation of the chromosomes into a lampbrush configuration. The growing chicken oocyte model has a number of advantages: (1) giant oocyte nuclei can be rapidly isolated manually under a stereomicroscope without enzymatic digestion and with visual quality control of the nuclei (Krasikova *et al*. 2012); (2) the nucleolar organiser is inactivated in lampbrush stage oocytes of adult birds, i.e. depletion of ribosomal RNA is not required (Davidian *et al*. 2023); (3) the domestic chicken genome is completely sequenced and well annotated (Huang *et al*. 2023); (4) the accumulated data on gene expression during oogenesis and early embryogenesis of the domestic chicken allow comparative functional analysis. Maternal RNA accumulates in the avian oocyte during the lampbrush stage of oogenesis, with RNA levels comparable to those found in the *Xenopus* oocyte (Olszanska and Stepinska 2008). It is also suggested that zygotic genome activation during chicken embryogenesis occurs in two waves, the first involving only maternal genes (Hwang *et al*. 2018b). At the same time, the transcriptome profile of avian oocytes has so far only been studied in early, primordial follicles (Rengaraj *et al*. 2014; Hu *et al*. 2020), in whole late follicles (Wang *et al*. 2017, 2018; Peng *et al*. 2019; Nie *et al*. 2022), or in the whole ovaries (Yin *et al*. 2020). Despite the rather detailed examples of nascent transcripts from tandem repeats (Trofimova and Krasikova 2016), the set of genes transcribed on the lateral loops of avian lampbrush chromosomes remained unknown.

Here, we aimed to determine a full spectrum of sequences transcribed on the lateral loops of chicken lampbrush chromosomes. To this end, we have systematically characterised the transcriptome in the nuclei and cytoplasm of chicken oocytes at the lampbrush chromosome stage of oogenesis. We examined nuclear and cytoplasmic mRNAs, as well as long and small non-coding RNAs. We also compared the obtained transcriptomic data with the available data on the transcriptomes of early chicken embryos. The results of this study revealed the contribution of transcripts synthesised on the lateral loops of lampbrush chromosomes to the pool of maternal RNAs, which play an essential role in the earliest stages of embryogenesis. Furthermore, we argue that hypertranscription of maternal mRNA and non-coding RNA genes defines the pattern of transcription loops along chicken lampbrush chromosomes.

## 2. Materials and Methods

### Preparation of nuclear and cytoplasmic RNA samples from chicken diplotene oocytes

Ovaries were taken from adult egg-laying chicken (*Gallus gallus domesticus*) females immediately after euthanasia. The experimental procedure was approved by the Ethics Committee for Animal Research of St. Petersburg State University (protocol #131-04-6 dated 25.03.2019). Unlike amphibians, growing oocytes in bird ovaries are at different stages of follicle maturation (Nie *et al*. 2022). Under the stereomicroscope, we manually separated pre-hierarchical follicles: small white follicles, 0.5-2 mm in diameter, containing oocytes at the lampbrush chromosome stage, and large white follicles, 2-4 mm in diameter, containing oocytes at the post-lampbrush chromosome stage. Next, we manually dissected nuclei from the selected oocytes using a Leica S9D or M165C stereomicroscope (Leica Microsystems) in 5:1 medium (83.0 мМ KCl, 17.0 mM NaCl, 6.5 mM Na2HPO4, 3.5 mM KH2PO4, 1 mM MgCl2, 1 mM dithiothreitol) as previously described (Krasikova *et al*. 2012). Additional rounds of washing of the isolated nuclei in 10 to 15 changes of 5:1 medium were carried out. Visual inspection of isolated nuclei under ×5-8 magnification confirmed that they contained fully developed lampbrush chromosomes (from oocytes of 0.5-2 mm in diameter) or partially condensed post-lampbrush chromosomes (from oocytes of 2-4 mm in diameter). Intact oocyte nuclei without nucleoplasm leakage were transferred one each into 5:1 medium and used for RNA extraction. Oocyte cytoplasm (ooplasm) was collected from enucleated oocytes to avoid contamination with nuclear RNA (Table S1).

### RNA extraction and RNase R treatment

Total RNA, including small RNA, was isolated from the nuclei and cytoplasm of growing chicken oocytes using TRIzol reagent (ThermoFisher Scientific) according to the manufacturer’s recommendations, following a previously developed protocol (Krasikova and Fedorov 2016). RNA integrity was assessed using RNA 6000 Pico Kit for Bioanalyzer (Agilent Technologies, #5067-1513). RIN > 7 in cytomplasmic samples confirmed good quality of isolated RNA. RIN value was not applicable to chicken oocyte nuclear RNA samples due to inactivation of nucleolar rRNA genes (Krasikova and Fedorov 2016).

For RNase R treatment, the nuclear RNA precipitate was dissolved in 14 μl RNase-free water, denatured at 65°C for 5 min and split into two samples. RNAse R buffer (10×) and 40U RNasin (Silex) were added to both samples; 20U (1 μl) RNase R (Epicentre) or 1 μl RNase-free water were added to experimental and control samples, respectively. RNA samples were incubated for 1 hour at 37°C. RNA was then purified on affinity columns (RNA clean and concentrator-5 kit, Zymo Research) and finally eluted in 6 μl of RNAse-free water.

### Sequencing of long and small RNAs

RNA concentration was measured using Qubit™ RNA HS Assay Kit (ThermoFisher Scientific, #Q32852). Libraries from oocyte cytoplasm RNA fractions were prepared with a starting amount of 100 ng and amplified with 15 PCR cycles. For samples from the nucleus fraction the concentration of RNA was too low to quantify it with the Qubit but was still enough to generate libraries with an amplification step being 15 PCR cycles for long RNA libraries and 25 cycles for small RNA libraries.

#### Long RNA libraries

For poly(A) RNA libraries, mRNA was isolated using NEBNext Poly(A) mRNA Magnetic Isolation Module (New England BioLabs, #E7490). Poly(A) RNA libraries were constructed using NEBNext Ultra II RNA Library Prep Kit for Illumina (New England BioLabs, #E7770) following manufacturer’s instructions. Total RNA libraries were constructed using NEBNext Ultra II Directional RNA Library Prep Kit for Illumina (New England BioLabs, #E7760) following manufacturer’s instructions. RNA was fragmented to the average size of 200-300 nt by incubating at 94°C for 5-10 min. Libraries were quantified with Qubit dsDNA HS Kit (ThermoFisher Scientific #Q32854) and the size distribution was confirmed with High Sensitivity DNA Kit for Bioanalyzer (Agilent Technologies #5067-4626). Poly(A) RNA libraries were sequenced on Illumina NextSeq 500 in single read mode with read length 76 bp following manufacturer’s instructions. Total RNA libraries were sequenced on Illumina HiSeq 4000 in paired-end mode with read length 2 x 151 bp following manufacturer’s instructions.

#### Small RNA library

Libraries of small RNA were constructed using the NEBNext Multiplex Small RNA Library Prep Set for Illumina (New England BioLabs, #E7330) following manufacturer’s recommendations. Purification of nucleotide fragments was conducted using high-resolution gel containing 0.8% agarose, 0.4% polygalactomannan and 1.6% γ-polygalactomannan in TAE buffer. Gel slices corresponding to fragments approximately 147 - 160 bp in size were then extracted using the MinElute Gel Extraction Kit (Qiagen, #28604). Libraries were quantified with Qubit dsDNA HS Kit (ThermoFisher Scientific, #Q32854) and the size distribution was confirmed with High Sensitivity DNA Kit for Bioanalyzer (Agilent Technologies #5067-4626). Libraries were sequenced on Illumina NextSeq 500 in single read mode with read length 76 bp following manufacturer’s instructions.

### RNA-seq data analysis

#### Trimming of reads

Reads were trimmed by Fastp 0.21.0 (Chen 2023) using the following techniques. For single-end libraries, adapters were trimmed using the adapter sequence AGATCGGAAGAGCACACGTCTGAACTCCAGTCA. For paired-end libraries, adapter sequences were determined automatically by read overlap analysis and then trimmed. Bases with Phred quality score below 3 were trimmed from 3’ ends of reads. If a region of 5 consecutive bases had an average Phred quality score below 15, these bases and everything towards the 3’ end of the read were trimmed. If after the above-mentioned procedures a read had an average Phred score below 20, this read was entirely discarded. If the discarded read belonged to a paired-end library, its mate was discarded too.

#### Read alignment and coverage analysis

Reads of long RNA libraries were aligned to the latest chicken genome assemblies - GRCg6a or telomere-to-telomere chromosome assembly GGswu1 (Huang *et al*. 2023) by STAR-2.7.3a (Dobin *et al*. 2013) with the option “--genomeSAindexNbases 14”. Reads of small RNA libraries were aligned to GRCg6a or GGswu1 genome assemblies by Bowtie 1.3.0 (Langmead *et al*. 2009) using the following options: “-n 1 −l 10 -k 1 --best”.

RNA coverage tracks and annotated genes with the exon-intron structure were visualised using the Integrative Genomics Viewer (IGV) (Robinson *et al*. 2011). Repetitive element positions were localised using the Censor tool with the chicken repeat library from Repbase (Kohany *et al*. 2006).

To reveal the expressed genes, the RNA-seq data was summarised as FPKM (Fragments Per Kilobase of transcript per Million reads mapped) values by the Cufflinks programme (Version 2.2.1) (Trapnell *et al*. 2012), followed by zFPKM transformation (Hart *et al*. 2013). According to (Hart *et al*. 2013), we consider genes with zFPKM ≥ −3 as expressed.

#### Classification of small RNAs

To remove possible contamination, reads of small RNA libraries were aligned to GGswu1 assembly by BBMap 39.01 (https://sourceforge.net/projects/bbmap/), with the word size of 8 bp and the minimum sequence similarity of 93.3%. Reads that did not align were not used in further analyses.

To accelerate and simplify subsequent operations, reads were dereplicated by VSEARCH 2.22.1 (Rognes *et al*. 2016). During dereplication, all identical reads were replaced with one read, keeping information about copy number; then all reads with copy number below 3 or length below 15 bp were discarded.

To classify small RNAs, the reads were aligned to a FASTA file that was constructed by combining the following sequences: (i) sequences of non-coding RNAs of *G. gallus* from Release 106 of ENSEMBL (Cunningham *et al*. 2022), (ii) sequences of mature miRNAs of *G. gallus* from miRBase (Kozomara *et al*. 2019), (iii) sequences of piRNAs of *G. gallus* from piRBase (Wang *et al*. 2022), (iv) sequences of nuclear tRNAs of *G. gallus* from genomic tRNA database (Chan and Lowe 2009). The above-mentioned databases were current as of 20/06/2023.

The alignment of reads was performed by BLASTN 2.11.0 (Camacho *et al*. 2009) with a word size of 7 bp. No restrictions for e-value were used. Only the best matches were considered. Matches were ignored for reads shorter than 40 bp if they had an edit distance greater than 1 and for reads longer than or equal to 40 bp if they had an edit distance greater than 2. The edit distances were calculated relative to the shorter sequence.

Reads whose matches did not pass the thresholds were considered to be piRNA if their length was 24-31 bp and siRNA if their length was 21-22 bp. Also, if the best match of a read was to a reference lncRNA and the read length was 24-31 bp, the read was considered to belong to a piRNA.

The above method allowed classifying reads into the following categories based on their best matches or lengths: miRNAs, siRNAs, piRNAs, snRNAs, snoRNAs, scaRNA, ribozyme, lncRNA, tRNA, rRNA, others. ‘Others’ is the category for reads that had no matches and at the same time their lengths did not fall within the 21-22 bp (siRNA length range) and 24-31 bp (piRNA length range) ranges.

#### Classification of targets of piRNAs

To determine piRNA targets, we annotated repeats in GGswu1 genome assembly. Repeat families were determined by RepeatModeler 2.0.3 (Flynn *et al*. 2020) with RMBLAST as the alignment tool (the option “-engine rmblast”) and with advanced detection of LTR transposable elements (the option “-LTRStruct”). Locations of repeats belonging to the found repeat families were determined by RepeatMasker 4.1.2 (Smit *et al*. 2013) with option “-e rmblast”.

piRNA sequences, determined as described in the “Classification of small RNAs” section, were aligned by BLAST to the repeats found by RepeatMasker. The alignment options and similarity thresholds were the same as described in the “Classification of small RNAs” section, except that instead of a single best match, 10000 best matches were considered. This allowed us to classify targets of piRNAs whose sequences aligned equally well with several different repeat families, using the same method as was used in (Pezic *et al*. 2014).

#### Strand-specificity of the transcripts

Reads were aligned against the GRCg6a assembly by HISAT (Version 2.2.1) (Kim *et al*. 2019) and then transcripts were identified by StringTie (Version 2.2.1) with the minimum allowed transcript length 200 nucleotides (Pertea *et al*. 2015). To search for overlapping transcripts from the opposite strands we used an in-house workflow. Strand-specific RNA-seq data was visualised with R package Gviz using custom import function strandedBamImport.

### Circular RNA identification

The programme “fast_circ.py” with the “de novo” option that comes with CIRCexplorer2 (Version 2.3.8) (Ma *et al*. 2021) was used with default parameters for *de novo* prediction of circular RNAs.

### Enrichment analysis and transcriptomic data visualisation

Phantasus software was used to perform k-means clustering and heatmap visualisation (https://ctlab.itmo.ru/phantasus/) (Kleverov *et al*. 2022). Gene set enrichment analysis to find over-represented (P < 0.001) gene ontology (GO) terms, key transcription regulators and histone modifications was performed using ENRICHR software (http://amp.pharm.mssm.edu/Enrichr) (Kuleshov *et al*. 2016). Database miRTarBase (https://mirtarbase.cuhk.edu.cn/) (Huang *et al*. 2022) was used as a source of experimentally validated miRNA targets. Visualisations were done with R software, version 4.2.2. For clustering analysis and visualisation purposes RNA-seq data were summarised as TPM (Transcripts Per Million) values.

Tissue-specificity index (tau) was used to measure tissue-specificity of gene expression (Yanai *et al*. 2005). Tau index ranges from 0 (broad expression) to 1 (narrow expression). A gene was considered as tissue-specific for Tau ≥ 0.95 as previously proposed (Jehl *et al*. 2020). Tau values for genes expressed across chicken organs were obtained from (Bush *et al*. 2018).

### RNA directed fluorescence *in situ* hybridisation on isolated lampbrush chromosomes

FISH probes were prepared from the chicken BAC-clone library CHORI-261 (https://bacpacresources.org/chicken261.htm) to the selected genomic regions (Table S2). BAC-clone based probes were labelled either with biotin-11-dUTP (Lumiprobe), digoxigenin-11-dUTP (Jena Bioscience) or aminoallyl-dUTP-ATTO-647N (Jena Bioscience) by nick-translation as described in (Krasikova *et al*. 2023a). To visualise clusters of CNM repeat together with the centromere repeat of the chromosome 3 (Cen3), we performed FISH with Cy3-labelled CNM-neg oligonucleotide probe (Krasikova *et al*. 2006) followed by reFISH with biotinylated PCR-labelled Cen3 probe (Krasikova *et al*. 2012). Lampbrush chromosomes were isolated microsurgically from the nuclei of chicken diplotene oocytes under a stereomicroscope Leica S9D or M165C (Leica Microsystems) according to the earlier described protocol (https://projects.exeter.ac.uk/lampbrush/protocols.htm). Special attention was given to RNA preservation during chromosome isolation. FISH on lampbrush chromosome preparations was performed according to DNA/RNA- and DNA/DNA+RNA-hybridisation procedures to visualise gene transcripts and non-transcribed genomic regions as detailed in (Zlotina and Krasikova 2017; Kulikova and Krasikova 2022). Hybridisation mixtures (20-30 ng/µl of BAC-clone DNA probe, 50% formamide, 50× excess of salmon sperm DNA, 10% dextran sulfate and 2×SSC) were denatured at 95°C for 10 min. Following the DNA/DNA+RNA hybridisation protocol, chromosome preparations were denatured in 70% formamide in 2×SSC at 70°C for 10 min. A combination of control experiments was performed to assess the quality of spreads, RNA integrity and appearance of nascent gene transcripts on lampbrush chromosome preparations (Kulikova and Krasikova 2022).

### Microscopy and image analysis

Chromosomes were analysed using a Leica DM4000 epifluorescence microscope (Leica Microsystems), with images captured by a CCD camera (1.3 Mp resolution). DAPI-stained chromosomes were imaged prior to FISH experiments. To ensure the accuracy of FISH mapping, at least 5 images from different chromosome sets were acquired. For each mapped genomic region, schematic drawings representing labelled lampbrush chromatin domains were generated. Contour length of transcription loops was measured using the Curved Line tool in ImageJ.

### Data access

RNA-seq data is available at the NCBI Sequence Read Archive (SRA) via accession numbers provided in the Supplementary Table S1.

## 3. Results

### Cytoplasmic and nuclear RNA fractions from chicken oocytes

In the ovaries of sexually mature chickens, the bulk of the oocytes remain in the diplotene stage of the first prophase of meiosis throughout the entire period of egg production. We isolated RNA from the lampbrush stage oocyte cytoplasm and nuclei manually dissected under a stereomicroscope (Figure 1 a; Table S1). Consistent with our previous findings (Krasikova and Fedorov 2016), 18S and 28S ribosomal RNA (rRNAs) peaks and both short and long RNAs were detected in the oocyte cytoplasm, whereas no nucleolar rRNA was observed in the oocyte nuclei, which were instead enriched in low molecular weight RNAs (Figure 1 b-b’, c-c’).

**Figure 1.**
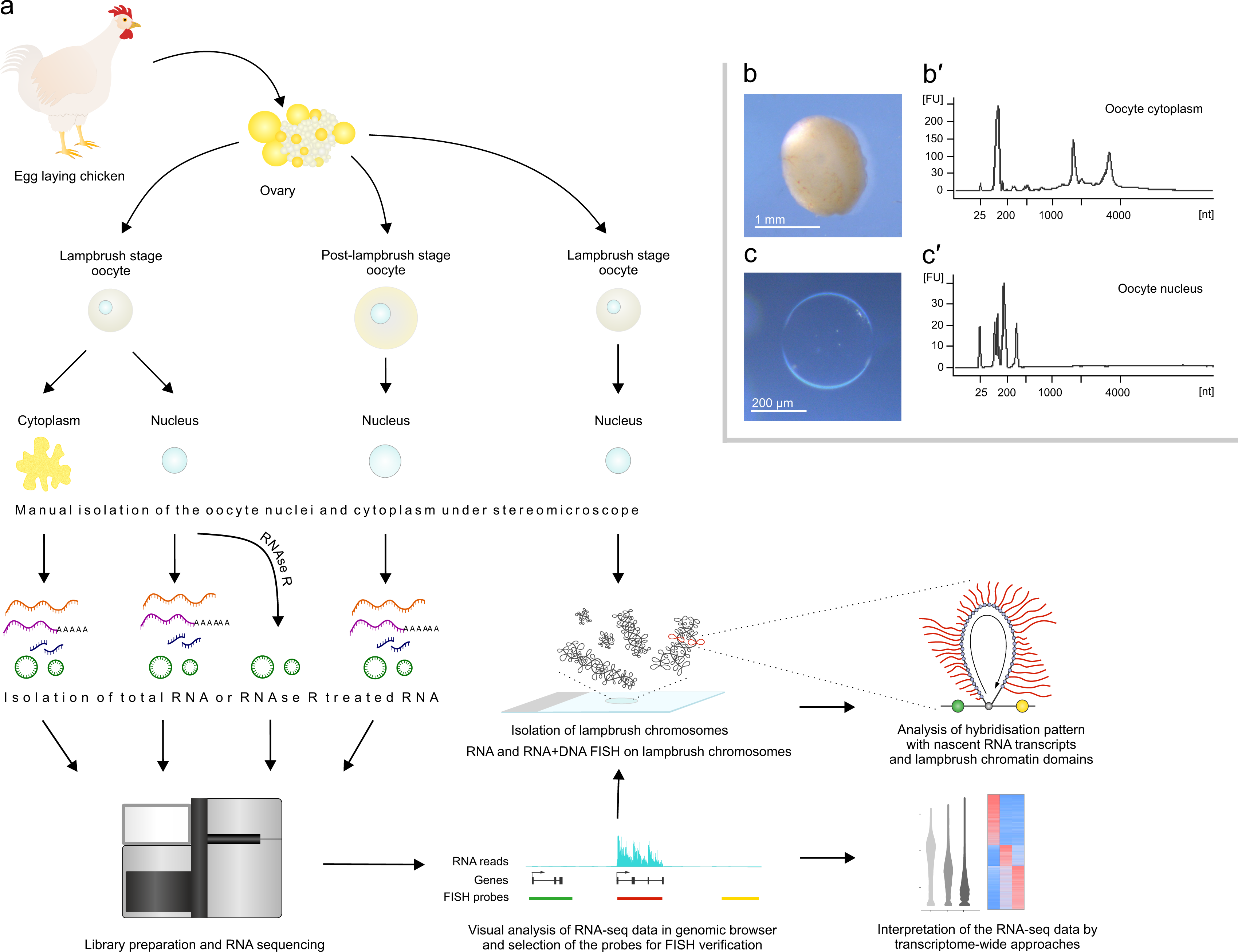
Chicken oocyte nucleus whole transcriptomic profile defines the pattern of genes transcribed by the lampbrush chromosomes. **a.** Experimental design for genome-wide transcriptome analysis of the chicken oocyte nuclear and cytoplasmic RNA combined with visualisation of nascent RNA transcripts and lampbrush chromatin domains on isolated lampbrush chromosome preparations. **b, b’, c, c’.** Cytoplasmic and nuclear RNA fractions from chicken diplotene oocytes. **b** - small white follicle containing oocyte at the lampbrush chromosome stage. **c** - manually dissected intact oocyte nucleus (or germinal vesicle) without nucleoplasm leakage. **b’**, **c’** - total RNA profiles for isolated oocyte nuclei (**c’**) and cytoplasm from enucleated oocytes (ooplasm) (**b’**). 18S and 28S ribosomal RNAs (rRNAs) peaks were detected only in the oocyte cytoplasm.

To obtain a complete picture of the primary RNA molecules, we performed strand-specific total RNA-seq of nuclear and cytoplasmic RNA fractions from lampbrush-stage oocytes. The nuclear RNA was devoid of transcripts from mitochondrial genes and the cytoplasmic RNA was devoid of transcripts from erythrocyte-specific genes, demonstrating that pure cytoplasmic and nuclear RNA fractions can be prepared from chicken oocytes. For the study of polyadenylation we sequenced poly(A)-enriched RNA libraries from oocyte nuclei and cytoplasm. For the study of small RNA species, we performed high-throughput sequencing of small RNA fractions from both oocyte nuclei and cytoplasm (Table S1).

### Nuclear and cytoplasmic transcriptome profiles along the full length of telomere-to-telomere chicken chromosome assemblies

We first analysed the transcriptome profile along the full length of the chromosomes. Mapping of total RNA-seq reads against the complete sequence of the chicken genome (version GGswu1) is shown for each chromosome assembly, including sex chromosomes and dot chromosomes (Figure 2 a, Figure S1). We estimate that approximately 28% of the chicken telomere-to-telomere genome assembly is covered by RNA-seq reads with ≥10 read depth in the case of total RNA libraries from lampbrush stage oocyte nuclei (Figure S2).

**Figure 2.**
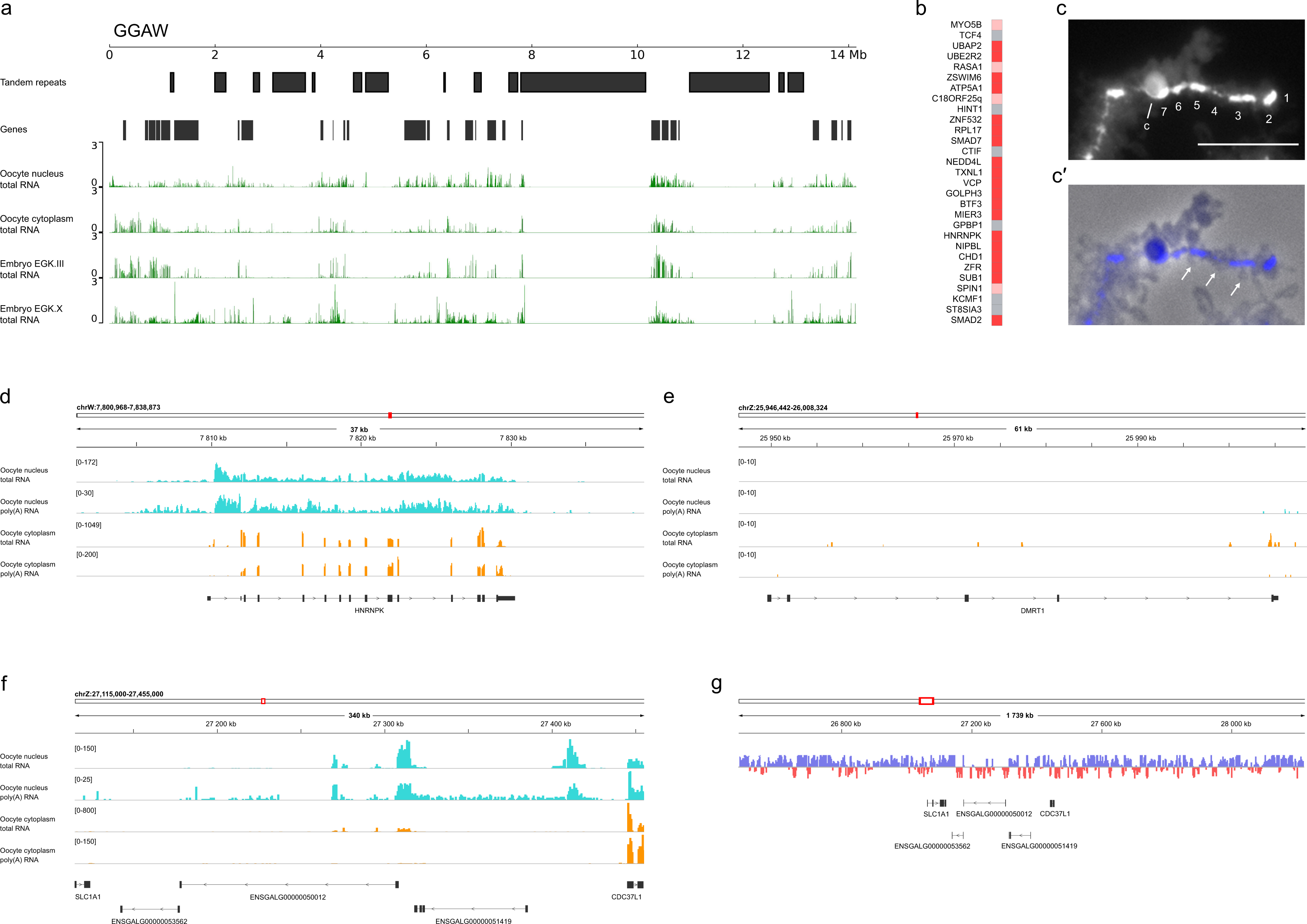
Sex chromosome gene expression at the lampbrush chromosome stage of oogenesis. a. W chromosome expression profile at the lampbrush chromosome stage of oogenesis. Total RNA profiles from chicken lampbrush-stage oocyte nucleus and cytoplasm against the chicken W chromosome assembly. Total RNA profiles for the EGK.III and EGK.X stages of embryogenesis along the W chromosome assembly are also shown. RNA-seq data for EGK.III and EGK.X stages of embryogenesis were obtained from (Hwang *et al*. 2018a). **b.** Expression levels of chicken W chromosome genes. Heatmap showing levels of transcripts for W chromosome genes in chicken oocyte nucleus at the lampbrush chromosome stage. Grey - no transcripts, pink - low level of transcripts, red - high level of transcripts. **c-c’**. Chromosome W at the lambrush stage, stained with DAPI (grayscale) (**c**) and merged with the phase contrast image (**c’**). Chromomeres are numbered from 1 to 7; arrows indicate small transcription loops; c - chiasmata. Scale bar – 10 μm. **d-f.** Examples of transcribed *HNRNPK* gene on GGAW (**d**), untranscribed *DMRT1* gene on GGAZ (**e**) and long non-coding RNA genes from the male hypermethylated region (MHM1) on GGAZ (**f**). Coverage tracks of total and poly(A) RNA from the oocyte nuclei (cyan) and cytoplasm (orange) of lampbrush stage oocytes along the gene body are shown; count range limits are given within square brackets. The exon-intron structure shown corresponds to one of the annotated transcript variants. **g.** MHM1 region is not hypermethylated in chicken lampbrush stage oocytes. Single-nucleus DNA methylation profile of chicken lampbrush chromosomes was obtained from (Nurislamov *et al*. 2022). For simplicity, only genes in the MHM1 locus are shown.

The higher density of reads, reflecting transcription in a particular genomic region, generally correlates with lampbrush chromosome segments with smaller chromomeres and longer lateral loops. For example, chromosome 3 in a lampbrush configuration has two main parts - the first with less compact, smaller chromomeres and longer lateral loops and the second with more globular, dense chromomeres with relatively short transcription loops (for example, Figure S1a in Krasikova *et al*. 2023b). We observe a higher density of transcribed sequences in the first part of lampbrush chromosome 3 (up to 57 Mb) compared to its second half (Figure S1). The density of RNA-seq peaks generally correlates with gene density along chromosomes, as does the density of expressed genes.

The W chromosome (GGAW), which is composed mainly of tandem repeats (more than 85% of the chromosome), at the lampbrush stage consists of 6-7 compact chromomeres (Solovei *et al*. 1998; Krasikova *et al*. 2006; Kulikova *et al*. 2020). Only a few tiny lateral loops can be seen between these large chromomeres (Figure 2 c, c’). We observe no or only a small number of RNA-seq reads in the genomic regions consisting of tandem repeats that correlates with the lampbrush configuration of the W chromosome (Figure 2 a).

Recently assembled heterochromatic dot chromosomes contain chromosome-specific multicopy genes that are expressed in testis (Huang *et al*. 2023). We analysed the total RNA-seq profiles along euchromatic and heterochromatic parts of the assembled dot chromosome models (GGA16, GGA29 to GGA32, GGA34 to GGA38) (Figure S1). In case of GGA29, which has the largest proportion of pericentromeric heterochromatin, we observe an almost complete absence of RNA reads aligned against the heterochromatic part in RNA samples from lampbrush-stage oocytes and early embryonic stages (Figure S1). This part of GGA29 contains 386 intact copies of an olfactory receptor *OR29G* gene (Huang *et al*. 2023). We assume that *OR29G* gene is not transcribed on chicken lampbrush chromosomes.

GGAW also contains multicopy gene *HINT1W* that is duplicated 52 times as a part of tandemly repetitive sequence (sate-HINT1) (Huang *et al*. 2023). We observed very low read coverage in the sate-HINT1 region in oocyte nuclear and cytoplasmic RNA-seq alignments (Figure 2 a, b).

### Oocyte cytoplasmic mRNA and long non-coding RNA consist primarily of spliced exons

Next we analysed gene expression profile for chicken oocytes at the lampbrush chromosome stage. In chicken oocyte cytoplasmic poly(A) RNA and total RNA fractions, we detected mature mRNAs and long non-coding RNAs consisting of spliced exons (Figure 3 a, b, Figure S3). The boundaries of the exons in mRNAs are typically sharply defined. In total, in chicken oocyte cytoplasm we detected spliced transcripts of 12312 genes (Supplementary Data Set 1). Similarly, in the *X. tropicalis* oocyte, cytoplasmic RNA consists primarily of spliced exons (Gardner *et al*. 2012).

**Figure 3.**
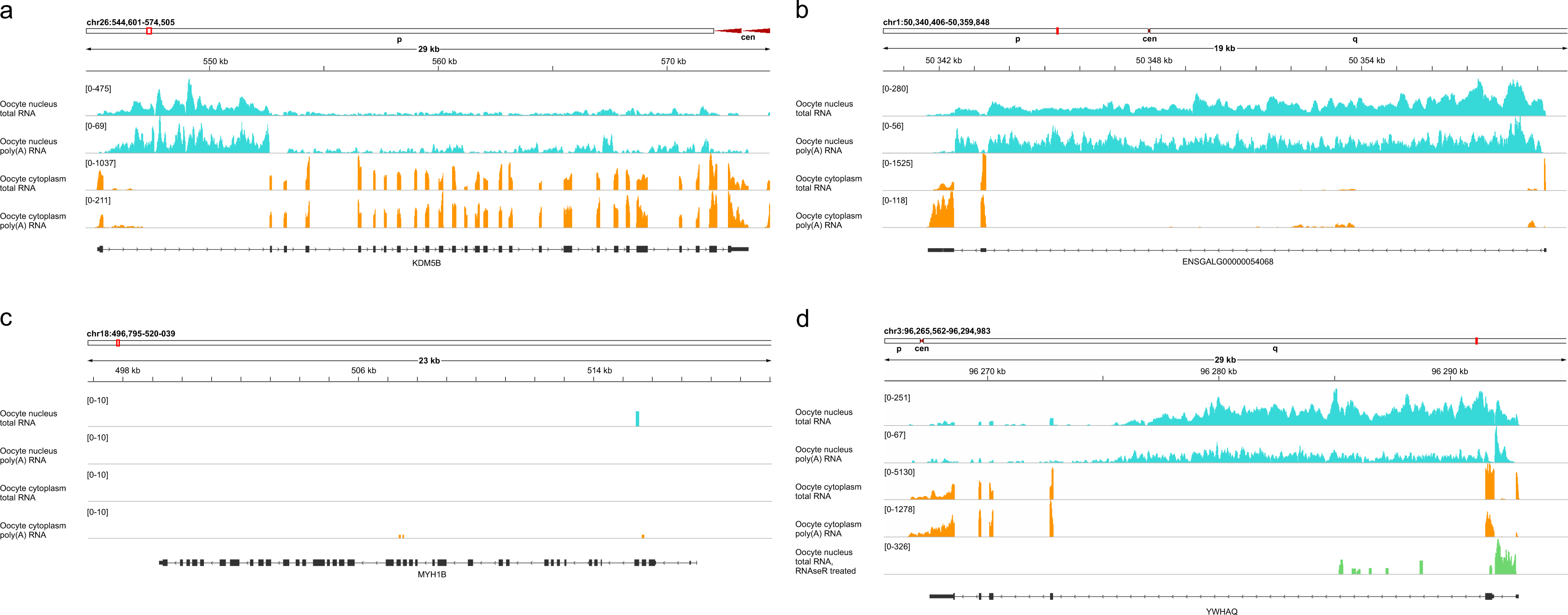
Nuclear and cytoplasmic total and poly(A) enriched RNA sequences from protein-coding and non-coding RNA genes. **a, b** - Oocyte cytoplasmic mRNA from *KDM5B* gene and long non-coding RNA from ENSGALG00000054068 gene consist primarily of spliced exons. Oocyte nuclear RNA contains nascent RNA and products of RNA processing for the same genes. **c** - Untranscribed protein-coding gene *MYH1B*. **d** - Stable intronic sequence RNAs (sisRNAs) resistant to RNase R treatment spliced from the nascent pre-mRNA transcript of *YWHAQ* gene. Coverage tracks of total and poly(A) RNA from the oocyte nuclei (cyan) and cytoplasm (orange) of lampbrush stage oocytes visualised by IGV browser along the gene body are shown; count range limits are given within square brackets. Coverage track of total RNA from the oocyte nuclei after RNase R treatment reveals sisRNAs (green). The exon-intron structure shown corresponds to one of the annotated transcript variants.

### Oocyte nucleus contains nascent RNA and products of RNA processing for the majority of expressed genes

In chicken oocyte nucleus we detected transcripts of 13185 genes (Supplementary Data Set 1). Most of these transcribed genes coincide with genes for which cytoplasmic spliced mRNAs or long non-coding RNAs were revealed. Visual examination of expression profiles for hundreds of these genes revealed that nuclear RNA contains both partially processed gene transcripts and intronic sequences with relatively low representation of exonic sequences (Figure 3 a, b, Figure S3). For poly(A) RNA libraries, the average ratio of exon and intron read coverage in the oocyte cytoplasm was 97.93, while in the nucleus it was only 1.38. Accordingly, in the case of total RNA from the oocyte cytoplasm, only ∼4% of the genome is covered by RNA-seq reads with ≥10 read depth (Figure S2).

In contrast to *X. tropicalis* oocyte nuclear RNA-seq profiles, in chicken oocyte nucleus we detected reads covering the entire length of the gene. This is due to the higher sequence depth of chicken oocyte nuclear RNA samples that contain no nucleolar rRNA, whereas *Xenopus* oocyte nucleus comprises hundreds of amplified nucleoli, and the rRNA was not removed before sequencing to reveal all types of RNA (Gardner *et al*. 2012). At the same time, we did not detect full-length unspliced primary gene transcripts for genes containing introns and exons. In fact, due to co-transcriptional splicing, we usually observe products of partial RNA processing even for nascent RNA (Beyer *et al*. 1981; Scherrer 2018). We observed partially processed transcripts with reads covering introns not only in total RNA libraries but also in poly(A) RNA libraries from the oocyte nucleus (Figure 3 a, b, Figure S3), since 3’-processing and polyadenylation of newly synthesised RNA can occur before splicing is complete (Lee *et al*. 2020).

Total RNA-seq of nuclear RNA fraction is more appropriate for nascent RNA detection. Moreover, the direction of transcription correlates with higher intron coverage at 5’ end (Gray *et al*. 2014). Here, on example of the genes with the lengthy introns we demonstrate the gradient in the distribution of intronic reads from 5’ to 3’ end (Figure S4).

Nuclear RNA sequencing leads to the identification of novel nuclear-retained long non-coding RNA species as transcription units that do not overlap with annotated genes (Mitchell *et al*. 2012). Similarly, we observed examples of spliced and polyadenylated unannotated transcripts in nuclear RNA fractions that can be considered as novel long non-coding RNAs, usually masked by more abundant cytoplasmic mRNAs (for example, Figure 9 f).

### The majority of genomic regions are transcribed from one strand

In our stranded nuclear RNA-seq data, we observed only few examples of genomic regions being transcribed from both strands, implying that the majority of genomic regions are transcribed from one strand (Figure S5 a-c; Supplementary Data Set 2). In the very few examples of transcripts from overlapping genes found in the oocyte cytoplasm, the transcript from one strand is derived from intranuclear nascent transcript, while the transcript from the other strand is observed only in the cytoplasm (Figure S5 d). We suggest that in such rare cases, the cytoplasmic transcript may be transcribed at an earlier stage of oogenesis.

### Most of gene transcripts are initiated at promoters, normally terminated and polyadenylated

In the vast majority of lateral loops on chicken lampbrush chromosomes, transcription is initiated at 5’ ends of genes coinciding with promoters (Figure 3, Figure S3, Figure S5). Our data also indicate that most gene transcripts are normally terminated, and we observed only rare examples of read-through transcription.

Polyadenylation of the transcripts regulates RNA stability during development and extends the maternal mRNA lifespan (Liudkovska and Dziembowski 2021). By sequencing poly(A)-enriched RNA libraries from the oocyte nucleus and enucleated cytoplasm we demonstrated that the majority of mRNAs and long non-coding RNAs in chicken oocyte are polyadenylated (Figure 3, Figure S3). On the RNA-seq profiles, the total RNA reads span to the termination of transcription while poly(A) RNA reads span till the site of polyadenylation. As visualised by scatter plots, the levels of gene transcripts in the polyadenylated RNA and total RNA libraries show a positive correlation both in oocyte nucleus (r = 0.81, p-value < 2.2e-16) and oocyte cytoplasm (r = 0.94, p-value < 2.2e-16) (Figure S6). Some gene transcripts are much more abundant in total RNA libraries than in polyadenylated RNA libraries, including histone genes, because their mRNAs are not polyadenylated (Marzluff and Koreski 2017). A number of long non-coding RNAs without poly(A) tails are found in the nucleus (Figure S6 a).

To examine dynamics of RNA synthesis during oocyte growth, we sequenced total nuclear RNA from late-stage oocytes, when chromosome condensation begins and lateral loops become shorter. Visual inspection of the RNA-seq profiles revealed that even in the late-stage oocyte nuclei, reads aligning the entire transcription units of the transcribed genes, from the transcription start site to the site of transcription termination, can be observed (Figure S4). The gradient of read coverage along an intron can be used to estimate the number and the speed of elongating RNA polymerases at gene body (Gray *et al*. 2014). Consistently, the gradient in read coverage for the intronic regions was less evident in the total RNA-seq data from oocytes at post-lampbrush chromosome stage (Figure S4).

### Cytoplasmic and nuclear transcripts emerge from nascent transcripts on the lateral loops of chicken lampbrush chromosomes

Since we detected unspliced or partially spliced nuclear transcripts for many genes, with reads covering the entire length of the transcription unit, we aimed to detect nascent transcripts on the transcription units *in situ* (Figure 1 a). Recently we developed an effective protocol that enables to map nascent gene transcripts on the lateral loops of lampbrush chromosomes by RNA-FISH (Kulikova and Krasikova 2022).

Here, by RNA-FISH-mapping with BAC-clone based probes we visualised transcription units of 9 genes at 8 chromosomal regions (Table S3). For all studied examples, we observed complete correspondence of transcript detection by nuclear RNA-seq and by RNA-FISH with nascent transcripts on lampbrush chromosome preparations (Figures 4, 5, Figure S7). These data are in line with previously reported correlation between nuclear RNA-seq and RNA-FISH estimations of nascent transcript levels (Mitchell *et al*. 2012).

**Figure 4.**
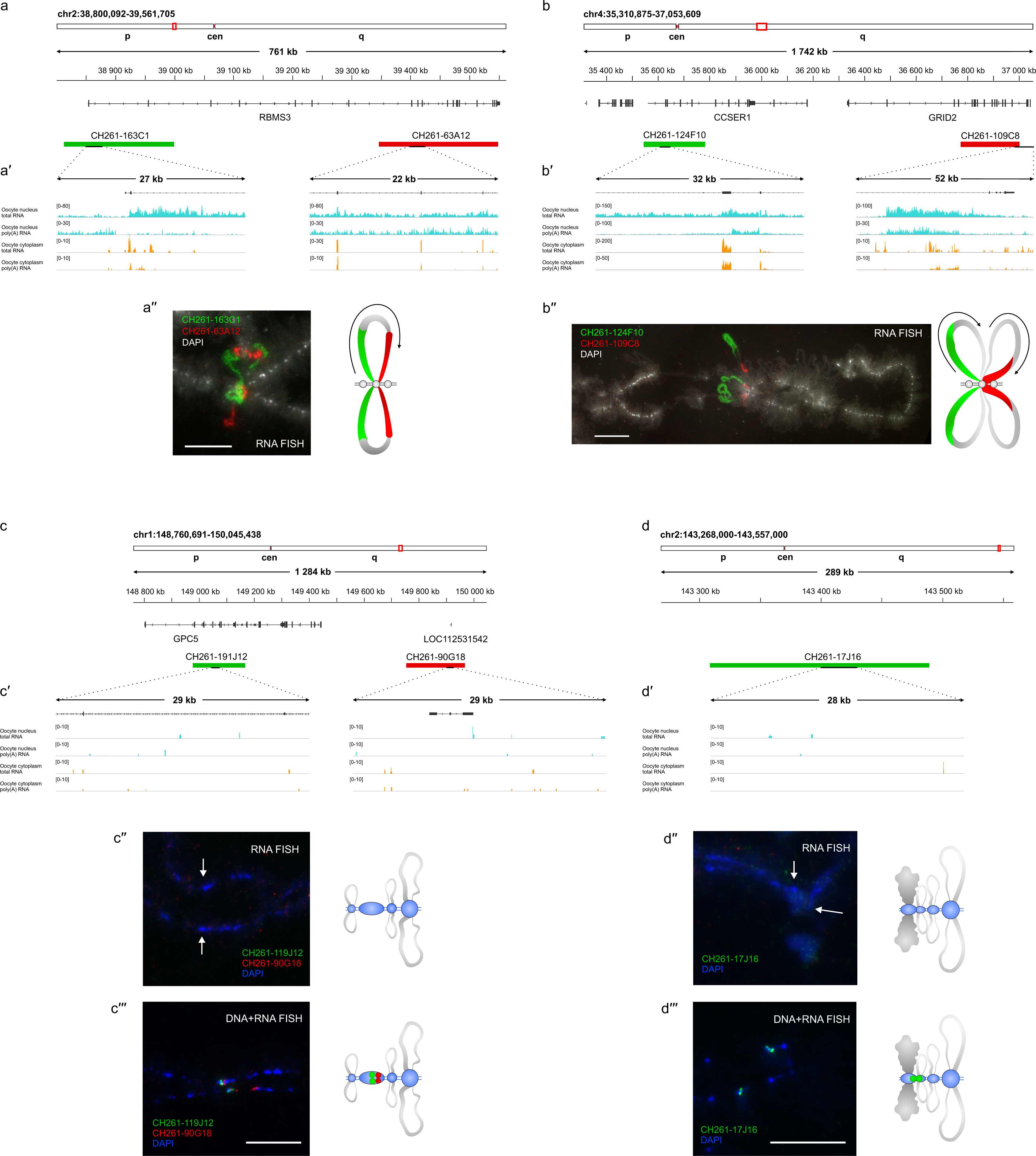
Visualisation of nascent pre-mRNA from individual transcribed protein-coding genes by RNA-FISH on lampbrush chromosomes. **a, b** - overview of the genomic regions containing transcribed genes *RBMS3* (a), *CCSER*1 and *GRID2* (b) with the positions of BAC clones used as FISH probes. **c, d** - overview of the non-transcribed genomic regions on chromosome 1 (c) and 2 (d) with the positions of BAC clones used as FISH probes. **a′, b′, c′, d′** - coverage tracks of total and poly(A) RNA from the nuclei (cyan) and cytoplasm (orange) of lampbrush stage oocytes visualised with IGV browser; data are shown for the short fragments covered by BAC clones; count range limits are given within square brackets. **a′′, b′′** - visualisation of nascent pre-mRNA from individual transcribed protein-coding genes. RNA-FISH with BAC clone-based probes CH261-163C1 (green) and CH261-63A12 (red) to the beginning and to the end of the *RBMS3* gene on chicken lampbrush chromosome 2 (**a′′**); RNA-FISH with BAC clone-based probes CH261-124F10 (green) to the beginning of *CCSER*1 gene and CH261-109C8 (red) to the end of *GRID2* gene on chicken lampbrush chromosome 4 (**b′′**); lateral loops are specifically labelled. **c′′, d′′** - visualisation of non-transcribed genomic regions falling into the chromomeres. RNA-FISH with BAC clone-based probes CH261-191J12 (green) and CH261-90G18 (red) on chicken lampbrush chromosome 1 (**c′′**) and BAC clone-based probe CH261-17J16 (green) on chicken lampbrush chromosome 2 (**d**′′); no signal can be detected. **c′′′, d′′′** - DNA+RNA-FISH using the same BAC clone-based probes as in panels c′′ and d′′; chromomeres, but not lateral loops, are specifically labelled. **a′′, b′′, c′′, d′′, c′′′, d′′′** - schematic drawings summarizing several FISH micrographs, arrows indicate the direction of transcription on the lateral loops. Scale bars – 10 μm. DAPI – greyscale (**a′′, b′′**) or blue (**c′′, d′′, c′′′, d′′′**).

Transcription pattern of the four sister chromatid copies of each gene were found to be identical in the RNA-FISH experiments (Figure 4 b’’). On the lateral loops we can see a gradient of tightly spaced nascent RNP fibers that coincides with the orientation of a transcribed gene and with the direction of transcription. Probes to the beginning of the gene can label the beginning of the transcription unit, whereas probes to the end of the gene label only the terminal part of the RNP matrix (Figure 4 a-a’’, b-b’’). At the post-lampbrush chromosome stage, the lateral loops transcribing the same genes become dramatically shorter, but we can still detect nascent transcripts (Figure S8).

We also conclude that the length of the transcription loops correlates with the length of the gene. One of the extremely long transcription units on lampbrush chromosome 14 (up to 57.7 μm in contour length) corresponds to the *RBFOX1* gene, which is 749 kb long and has intron sizes of 523-650 kb (Figure 5 a, b-b’’). Thus, the estimated compaction level of this gene is approximately 13 kb/μm, corresponding to a chromatin fibre that is only partially packed by nucleosomes. In contrast, transcription of the 77.1 kb *CSMD3* gene variant occurs on a much shorter lateral loop of approximately 3.3 μm in contour length (Figure S7 a-a’’).

**Figure 5.**
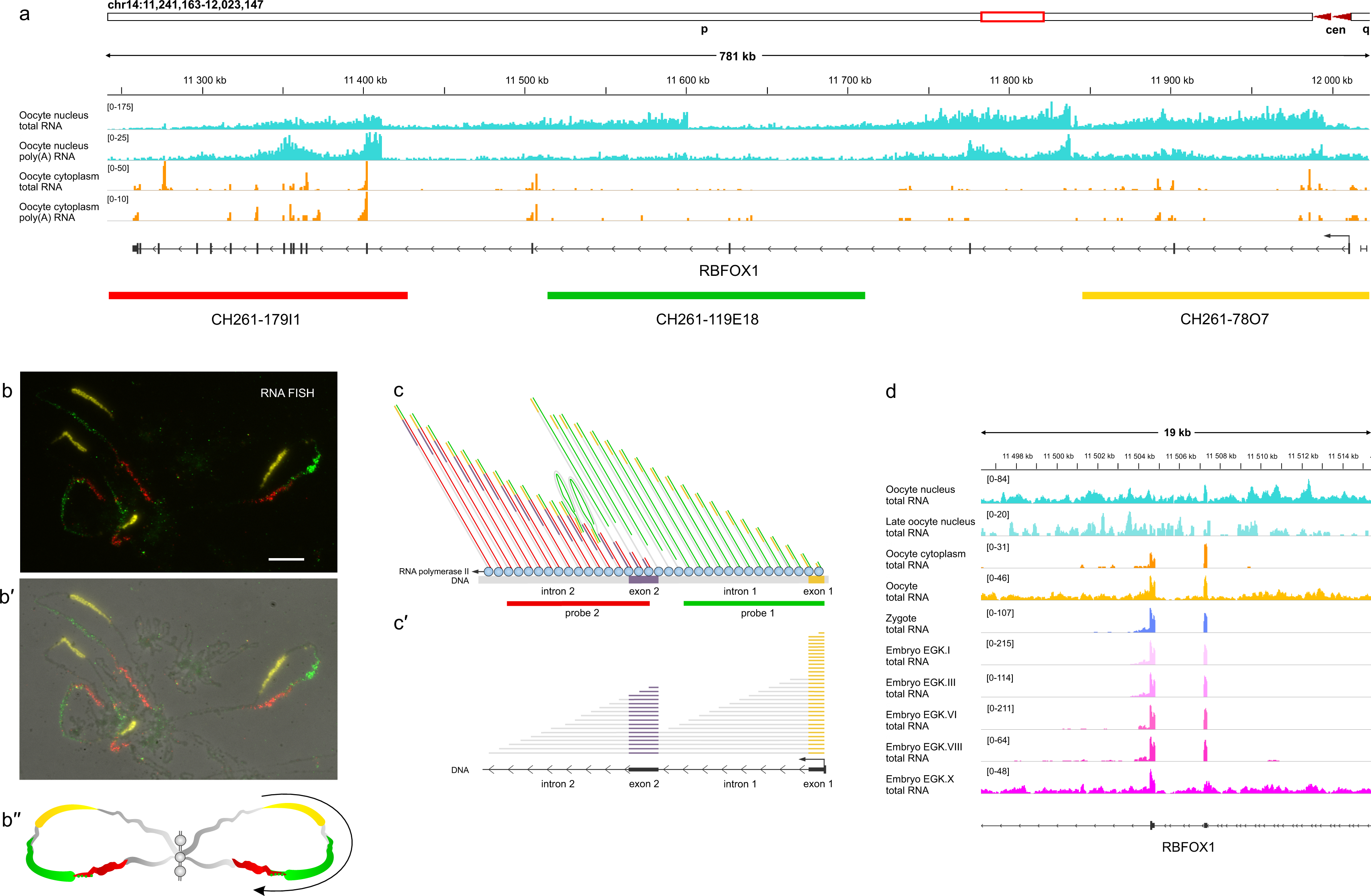
Maternal nascent pre-mRNA synthesis and co-transcriptional splicing on the lateral loops of lampbrush chromosomes on example of the *RBFOX1* gene with multiple lengthy introns. **a** - RNA-seq data of total and poly(A) RNA from the oocyte nuclei (cyan) and cytoplasm (orange) of lampbrush stage oocytes. Read coverage tracks visualised by IGV browser along the whole region of chicken chromosome 14 occupied by the *RBFOX1* gene are shown, count range limits are given within square brackets. The exon-intron structure shown corresponds to one of the transcript variants. The bottom row displays the positions of the BAC clones used as RNA-FISH probes, coloured as in **b-b′′. b-b′′** - RNA-FISH with BAC clones CH261-78O7 (yellow), CH261-119E18 (green), CH261-179I1 (red) to the beginning, to the middle and to the end of the *RBFOX1* gene on chicken lampbrush chromosome 14 (**b**), merged with the phase contrast image (**b′**) and schematic drawing summarising several RNA-FISH micrographs indicating direction of transcription (**b′′**). Scale bar - 10 μm. **c-c′** - diagram illustrating gradients in the length of nascent RNA visualised by RNA-FISH (**c**) and in the read coverage along an intron (**c′**) due to co-transcriptional splicing of nascent transcripts on example of a hypothetical gene. **d** - coverage track of RNA-seq data for lampbrush stage oocyte nucleus, post-lampbrush stage oocyte nucleus, oocyte cytoplasm, whole oocyte, zygote and embryonic stages EGK.I, III, VI, VIII, and X shown for a fragment (11,496,277-11,515,375 kb) of the *RBFOX1* gene with two exons, demonstrating preservation of the mature maternal *RBFOX1* mRNA from the oocyte to embryo stage EGK.VIII with the appearance of nascent zygotic transcripts at the EGK.X stage.

Another scenario - many genes of different polarity transcribed in a single lateral loop and separated by a chromatin knot (Figure 6) or even one by one without visible chromatin knots (Figure S9). Such arrangement of transcriptional units in lampbrush lateral loops completely corresponds to the classification of lateral loops in amphibian lampbrush chromosomes suggested according to electron-microscope observations of spread chromatin (Scheer *et al*. 1976).

**Figure 6.**
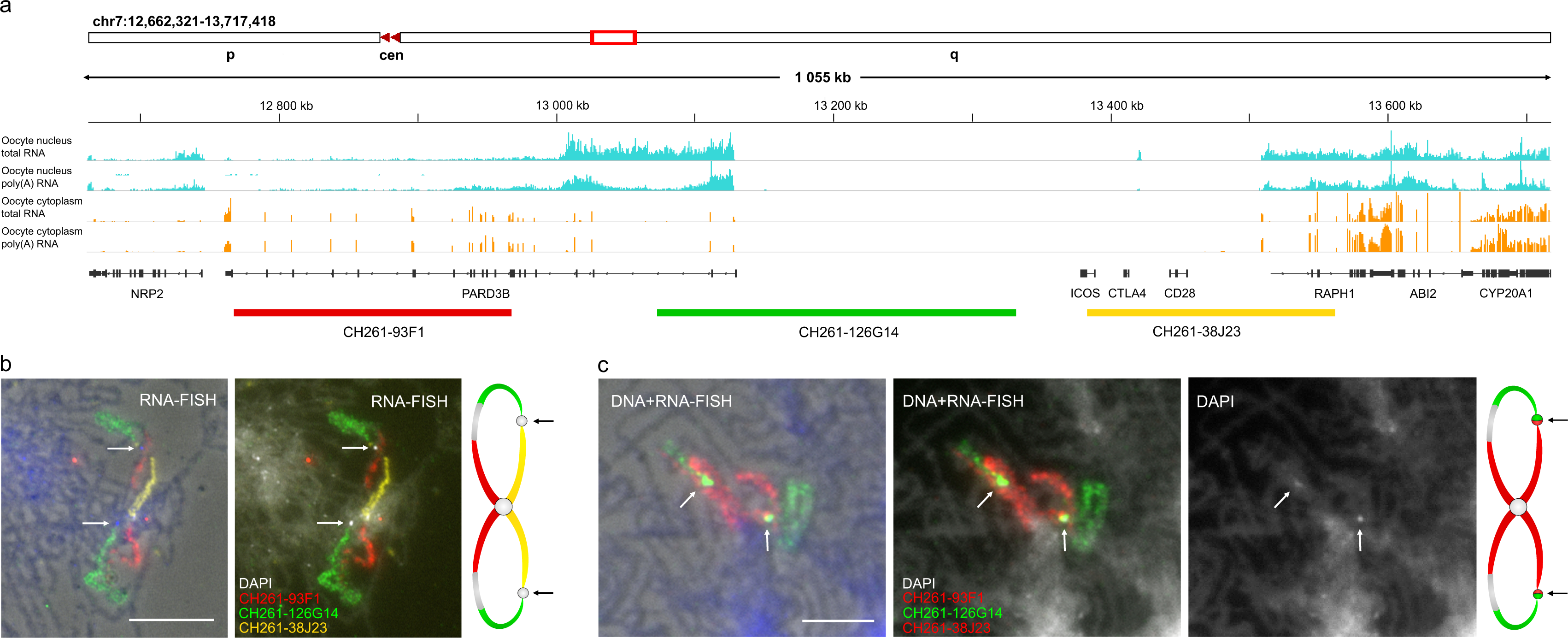
How the transcriptional activity is reflected in the chromatin structure on the example of the genomic region of chicken lampbrush chromosome 7. **a** - Genomic region on chicken chromosome 7 covered by BAC clones used as FISH probes (bottom row); coverage tracks of total and poly(A) RNA from the oocyte nuclei (cyan) and cytoplasm (orange) of lampbrush stage oocytes visualised with IGV browser. Count range limits are given within square brackets. A given chromosome segment contains transcriptionally active and silent genomic regions. **b**, **c** - RNA-FISH (**b**) and DNA+RNA-FISH (**c**) with BAC clones CH261-93F1 (red), CH261-126G14 (green) and CH261-38J23 (yellow or red) on chicken lampbrush chromosome 7; on the left panels fluorescent images are merged with the phase contrast images with DAPI staining shown in blue. RNA-FISH reveals transcribed genomic regions forming one lateral loop (**b**). DNA+RNA-FISH reveals chromatin knots on the axes of lateral loops, visible by DAPI staining (arrows) (**c**). Chromatin knot hybridizes with the parts of BAC clones CH261-126G14 (green) and CH261-38J23 (red) covering transcriptionally silent genomic regions (**c**). Schematic drawings summarise several RNA- and DNA+RNA-FISH micrographs. Scale bars – 10 μm. DAPI – grayscale or blue.

Although there is no doubt that thin-to-thick polarisation of the RNP matrix reflects the length gradient of nascent RNA and the direction of transcription (Gall 2014), our data suggest that such polarised units of RNP matrix with the same orientation within one lateral loop could correspond to a single gene with multiple lengthy introns. We noticed that in long transcription units gradients in nascent RNP matrix demonstrate cleavage of nascent transcripts, reproducing Christmas trees on the Miller spreads (Figure 5 b-b’’). Indeed, a drastic shortening of the nascent RNA can be observed after intron removal from the nascent transcript (Beyer *et al*. 1981). This pattern is reproduced by aligning nuclear RNA-seq data along the gene length, but with opposite orientation (Gray *et al*. 2014), and is shown schematically on example of a hypothetical gene (Figure 5 c-c’).

Three-coloured RNA-FISH with probes to the beginning, to the middle and to the end of the *RBFOX1* gene demonstrate co-transcriptional splicing of the nascent RNA on the lateral loops of the lampbrush chromosomes (Figure 5 a-b’’). Since we do not observe the overlapping of the signals from all three probes at the end of the transcription unit, intron removal occurred. Hybridisation signals from very short exonic regions in partially spliced nascent transcripts were not detectable in this transcription loop. Similar example is shown for the individual transcription loop for *RBMS3* gene whose nascent transcripts are detected by two probes to the beginning and to the end of the gene (Figure 4 a-a’’). No signal can be detected with the probe to the beginning of the *RBMS3* gene in the nascent transcripts at the end of the gene. These data additionally indicate that it would be hardly possible to sequence full-length pre-mRNA transcripts in the nuclear RNA samples, even when sequencing newly synthesised RNA.

Non-transcribed genomic regions according to RNA-seq data are found within chromomeres (Figure 4 c-c’’’; d-d’’’). For example, three probes to two genomic regions on GGA1 and GGA2 do not hybridise with nascent RNA in the RNA-FISH experiment (Figure 4 c’’, d’’), but label chromomeres after chromosomal DNA denaturation (Figure 4 c’’’, d’’’). Such genomic regions often contain no annotated genes (so-called gene deserts); they may also contain untranscribed genes. Interestingly, one lateral loop inserted into a chromomere can contain several individual transcription units separated by untranscribed chromatin knot or more rarely several knots (Figure 6). In some lampbrush chromosome preparations, such chromatin knots can rarely be found anchored to nearby chromomeres, leading to the formation of two smaller transcription loops.

### A number of nuclear intronic sequences are in a circular form

In the oocyte nuclear RNA certain intronic sequences were predominantly abundant within transcribed genes most probably reflecting separated intronic RNAs (Figure S3 g). High coverage of the individual introns in total RNA-seq of the nuclear RNA fraction appear as a single peak or as multiple peaks. Resistance to RNase R treatment indicates on the lariat form of some of the identified intronic sequences (Figure 3 d, Figure S3 g, h). Intronic origin RNAs thus represent stable intronic sequences RNA (sisRNAs), previously characterised in *X. tropicalis* oocyte nuclei (Gardner *et al*. 2012).

The appearance of mature mRNA for the sisRNA-producing genes suggests that they are spliced from the nascent transcript (Figure 3 d, Figure S3 g, h). Indeed, we checked that sisRNAs come from the same strand as pre-mRNAs. Furthermore, only certain intronic sequences are unusually stable, while the other introns from the same primary full-length transcript are degraded upon splicing. Some of the sisRNAs were observed as peaks not only in total RNA but also in the poly(A)-enriched fraction of the nuclear RNA. There is consistent evidence that hundreds of sisRNAs exist in a linear form in addition to a circular form (Chan and Pek 2019).

We next used CIRCexplorer2 (Ma *et al*. 2021) to identify circular RNAs (circRNAs) in total and poly(A)-enriched oocyte nuclear RNA samples and in total nuclear RNA sample after RNase R treatment. circRNAs were predicted *de novo* not only in RNA samples after RNase R treatment, but also in total and poly(A) RNA libraries; in the poly(A) RNA libraries they were represented by low read counts (Supplementary Data Set 3). We checked that the unmapped reads from the left side of the identified circRNA aligned with the sequence adjacent to the right side, representing chimeric reads. The list of the identified circRNAs and annotated genomic regions covered by each circRNA (including 5’UTR, exonic and intronic regions) is represented in Supplementary Data Set 3.

One of the examples of sisRNAs covering the 5’UTR is nuclear abundant RNase R resistant sisRNA of the *YWHAQ* gene (Figure 3 d). We suggest that 5’UTR-derived sisRNAs, either in linear or circular form, stored in the oocyte may be involved in the regulation of host gene expression.

### Overview of genes transcribed in the chicken lampbrush-stage oocyte

A total of 77% and 95% of all reads from the oocyte nuclear and cytoplasmic total RNA libraries, respectively, mapped within annotated genes. Similar to *Xenopus* oocyte (Gall, 2014), different genes vary greatly in the abundance of their cytoplasmic transcripts.

The chicken genome (version GRCg6a) contains 24356 annotated genes, including 16779 protein-coding genes and 5504 genes of long non-coding RNAs (lncRNAs). Transcripts for 10697 protein-coding genes and 2488 lncRNA genes in the oocyte nucleus and 11402 protein-coding genes and 910 lncRNA genes in the oocyte cytoplasm were detected (Supplementary Data Set 1).

### Comparison with chicken early embryo transcriptome suggests that oocyte nucleus is involved in maternal RNA synthesis

We further checked whether accumulated oocyte RNAs are stable until early stages of embryogenesis thus representing maternal RNA. In chicken there are two waves of zygotic genome activation: minor wave shortly after the fertilisation and major wave during cellularisation followed by maternal-to-zygotic transition (Rengaraj *et al*. 2020). Thus we compared the chicken oocyte transcriptome with RNA-seq data for Eyal-Giladi and Kochav stages I (EGK.I, early cleavage stage) and III (EGK.III, 80 to 90 cells) chicken embryos (Hwang *et al*. 2018a; b), before the second wave of zygotic genome activation occurs together with paternal genome activation (Rengaraj *et al*. 2020). For this purpose we have combined our transcriptome data with chicken embryos expression dataset (Hwang *et al*. 2018a).

Using the *RBFOX1* gene as an example, we show how maternal RNA transcribed on the lateral loops of lampbrush chromosomes is spliced into mature mRNA and stored in the oocyte cytoplasm before being transferred to the zygote and early embryos (Figure 5 d). Only at the EGK.X stage of embryonic development, transcription of the zygotic *RBFOX1* gene resumes, as evidenced by the appearance of RNA-seq reads along the intronic sequences. Comparative alignment of chicken oocyte cytoplasmic and nuclear total RNA-seq profiles as well as early embryonic stages EGK.III and EGK.X RNA-seq profiles (before and after the major wave of zygotic genome activation) along the telomere-to-telomere chicken chromosome assemblies is also shown on Figure S1. Note the very similar RNA-seq profiles between the oocyte cytoplasmic RNA fraction and the EGK.III embryonic RNA.

Next we described in more detail the expression of protein-coding genes and lncRNA genes in early embryos and oocytes. In total, transcripts of 64%, 68% and 65% of protein-coding genes and 45%, 17% and 16% of lncRNA genes were detected in the oocyte nucleus, oocyte cytoplasm and EGK.I embryo, respectively.

The sets of protein-coding genes whose transcripts are found in the oocyte nucleus, oocyte cytoplasm, and embryos overlap strongly (Figure 7 a). There is also a strong overlap between the sets of long non-coding RNA genes whose transcripts are found in the oocyte cytoplasm and in the embryos, with most of them also present in the oocyte nucleus (Figure 7 a). In addition, transcripts of a considerable number of long non-coding RNA genes are found only in the oocyte nucleus (Figure 7 a). The oocyte nucleus contains transcripts for more than 80% of the genes whose transcripts are found in the oocyte cytoplasm or early embryo. There is a positive correlation between the levels of gene transcripts in the oocyte nucleus and oocyte cytoplasm (r = 0.65, p-value < 2.2e-16) as well as in the oocyte cytoplasm and EGK.I embryo (r = 0.94, p-value < 2.2e-16) (Figure 7 b, c). Therefore, active transcription of these genes in the oocyte nucleus is the simplest and most likely mechanism to explain these observations.

**Figure 7.**
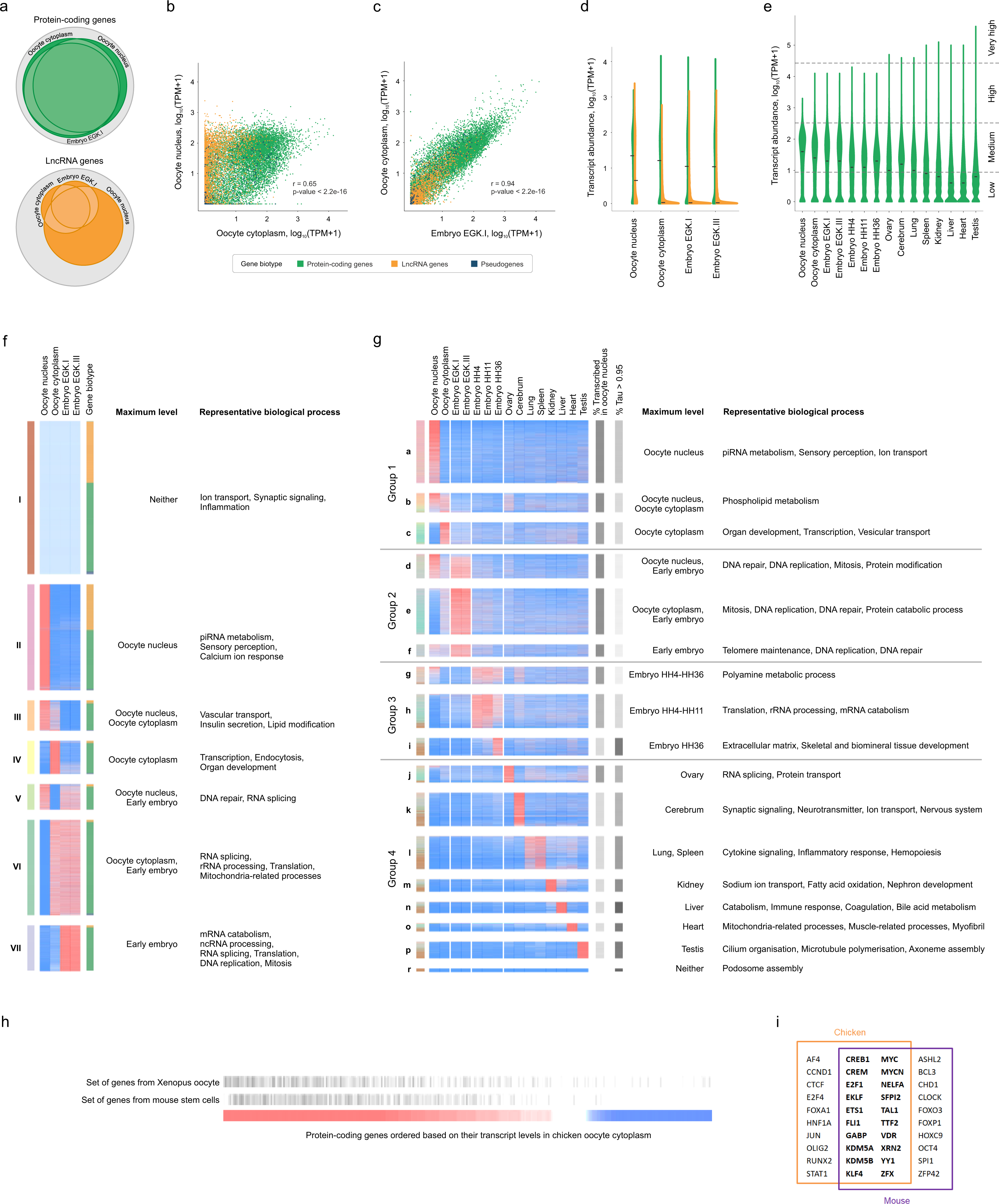
Comparison of transcriptomes and gene expression profiles in the oocyte nucleus, oocyte cytoplasm, early embryo and adult chicken organs. **a-c.** Euler diagrams (**a**) and scatter plots (**b, c**) illustrate the relationships between sets of protein-coding and long non-coding RNA (lncRNA) genes, whose transcripts were detected in the oocyte nucleus, cytoplasm and EGK.I embryo. On scatter plots each dot represents individual gene; colour of the dot corresponds to gene biotype. **d.** Violin plot illustrates the distribution of transcript abundance across expressed protein-coding and lncRNA genes in different oocyte compartments and early stages of embryogenesis. Chicken embryo expression dataset was obtained from the literature (Hwang *et al*. 2018a). **e.** Violin plot illustrates distribution of transcript abundance across expressed protein-coding genes in different oocyte compartments and adult chicken organs. Expression values were divided into 4 intervals: low, medium, high, and very high. Expression dataset for adult chicken organs was obtained from the literature (Bush *et al*. 2018). Black horizontal dashes in violin plots represent median values. **f.** Comparison of gene expression profiles in oocyte nucleus, cytoplasm and early stages of embryogenesis. Heatmap illustrates 7 clusters of co-expressed genes. Clusters are colour-coded as indicated by the column of coloured boxes on the left. The column of coloured boxes on the right indicates gene biotype: protein-coding genes (green), lncRNA genes (orange) and pseudogenes (blue). **g.** Comparison of gene expression profiles in oocyte nucleus, cytoplasm and adult chicken organs. Heatmap illustrates 17 clusters of co-expressed genes. The colours in the left column of coloured boxes correspond to the colours of the clusters shown on panel f. Two columns of boxes on the right indicate the proportion of genes with transcripts in the oocyte nucleus and the proportion of genes with narrow expression pattern (tau ≥ 0.95) for each gene cluster; minimum and maximum values are represented by white and black respectively. In both heatmaps, each row represents a gene and each column represents a sample. Expression values are visualised using a relative colour scheme, where maximum and minimum values in each row are represented by red and blue colours respectively. For each cluster of co-expressed genes main enriched representative biological process GO terms are listed. **h.** The transcriptional profile of chicken oocytes is similar to that of *Xenopus* oocytes and stem/progenitor cells of adult mouse organs. The gene set enrichment analysis plot shows the enrichment of the set of the top 3000 highly transcribed genes in the *Xenopus* oocyte (Jullien *et al*. 2014) and the set of the top 3000 genes that show strong correlations with stem/progenitor cell transcript levels in adult mouse organs (Kim *et al*. 2023) among the highly transcribed genes in the chicken oocyte. Protein-coding genes were ranked according to the level of their transcripts in chicken oocyte cytoplasm, high - red, low - blue. **i.** Comparison of sets of transcription regulators associated with hypertranscription. The first set (inside the orange box) represents transcriptional regulators associated with top 3000 genes demonstrating high transcript level in chicken oocyte cytoplasm. The second set (inside the purple box) represents transcriptional regulators associated with the top 3000 genes that show strong correlations with transcript levels in stem/progenitor cells in adult mouse organs (Kim *et al*. 2023). Both sets were produced by enrichment analysis for ChEA library of transcription factor targets. Results of enrichment analysis were ranked based on p-values. Top 30 unique transcription regulators from each set are shown. The two sets of transcription regulators share 20 out of the 30 members (shown in bold).

In the oocyte cytoplasm, the distribution of the RNA repertoire of both protein-coding genes and lncRNAs was similar to that of early embryos; the majority of lncRNAs had lower expression levels than protein-coding genes (Figure 7 d). In contrast, the distribution of the RNA repertoire of lncRNAs in the oocyte nucleus resembled that of protein-coding genes (Figure 7 d).

In order to perform a functional characterisation of genes expressed in the oocytes, we traced their expression level during early embryogenesis, identified seven clusters of co-expressed genes and found associated biological processes (Supplementary Data Set 4). Main enriched biological processes GO terms associated with clusters of co-expressed genes are indicated (Figure 7 f). For ∼50% of genes expressed in oocytes, expression was maintained or increased (clusters V, VI, VII) during early embryogenesis (Figure 7 f).

Enrichment analysis demonstrated that these genes are associated with essential cellular functions, including non-coding RNA processing, splicing, mRNA degradation, protein synthesis, mitochondria functions, DNA repair, DNA replication and cell cycle progression. It is noteworthy that genes whose transcripts are present at high levels in the oocyte cytoplasm and then decrease during early embryogenesis (cluster IV) are characterised by processes related to the regulation of transcription as well as developmental functions. lncRNAs are mainly found in cluster I of the non-expressed genes and in cluster II of the genes with the maximum accumulation in the oocyte nucleoplasm (Figure 7 f).

### Comparison with transcription profiles in adult organs: tissue-specificity of expressed genes

High-throughput analysis of the *Xenopus* oocyte transcriptome has demonstrated that “wide range of genes in lampbrush chromosomes accumulate a large number of transcripts” including “embryo-specific and cell-type–specific transcripts” (Simeoni *et al*. 2012). Thus we compared RNA repertoire in the oocyte nucleus and cytoplasm with the transcription profiles in different chicken organs. For this purpose we have combined our transcriptome data with chicken mRNA expression atlas (Bush *et al*. 2018).

Violin plot illustrates distribution of expression profiles of protein-coding genes in different oocyte compartments and chicken organs (Figure 7 e). The expression profiles were similar for the oocyte nucleus, cytoplasm and early embryos, where most of the transcribed genes showed medium levels of expression. In contrast, in differentiated tissues, most genes have low levels of expression, while a few genes have very high levels of expression (Figure 7 e). This difference in expression profiles is consistent with the assumption that differentiated cells from the organs express a narrow set of tissue-specific genes, whereas oocytes express a broader set of genes responsible for essential cellular functions.

In order to extend functional characterisation of genes expressed in the oocytes, we traced their expression in different chicken organs. Combined dataset was divided into 17 clusters of co-expressed genes using k-means clustering. Main enriched biological processes GO terms associated with clusters of co-expressed genes are indicated (Figure 7 g). Full lists of enriched biological processes GO terms associated with clusters of co-expressed genes (P < 0.001) are presented in Supplementary Data Set 5.

The identified co-expression clusters can be divided into 4 groups – those where the maximum expression is in oocytes (group 1), or in early embryos before zygotic genome activation (group 2), or in the embryos after genome activation (group 3), or in differentiated organs (group 4). Genes whose transcripts are present in the oocyte nucleus were distributed between groups 1, 2, 3 and 4 as follows: 34%, 26%, 16% and 24% of the genes, respectively. This may indicate that of the genes transcribed in the oocyte nucleus, the majority (76%) have a primary function in oogenesis or embryogenesis, while 24% also have functions in differentiated tissues.

For each cluster we calculated the proportion of genes whose transcripts are present in the oocyte nucleus (Figure 7 g). This parameter has the highest values in the clusters of groups 1 and 2 (> 90%) and in the ovarian-specific cluster (90%), decreases to 33% in the cluster for the embryo at post-ovipositional developmental stage HH36, and remains at the 40-60% level in all the clusters, except the ovarian, of group 4.

We then looked at the gene clusters, more than 90% of which are made up of genes transcribed in the oocyte (Figure 7 g). Among them is the cluster corresponding to the ovary (cluster j), which is not surprising. There are also clusters corresponding to the early embryo (before genome activation) (clusters e, f), indicating that the maternal transcripts accumulated in the oocyte are transferred to the embryo. There are also clusters corresponding to later embryos (after genome activation) (clusters g, h), suggesting that during the initial phase of genome activation, the same set of genes as in the oocyte is predominantly expressed. A notable difference between oocyte and embryo transcript profiles is observed at the HH36 embryo developmental stage (cluster i).

Next, for each cluster we calculated the proportion of genes with a narrow expression pattern (tau ≥ 0.95) (Figure 7 g). This parameter has the highest values in the clusters from the group 4 and the lowest values in the clusters specific to genes with maximum expression in the oocyte and, especially, in early embryos. Enrichment analysis revealed that clusters from the groups 1, 2 and 3 are mainly associated with essential cellular functions, whereas clusters from the group 4 are mainly associated with specific biological processes relevant to the physiology of the corresponding organ (Figure 7 g).

The data suggest that the set of genes transcribed in lampbrush stage oocytes is enriched in genes with a broad expression pattern (housekeeping genes). This may be because the oocyte and early embryo specifically require basic cellular processes to perform their functions.

### Key transcriptional regulators for genes expressed in lampbrush stage oocytes

To gain insight into the mechanisms of transcription regulation in chicken oocytes at the lampbrush stage, we checked whether the gene expression profile defined here is similar to that in other systems characterised by hypertranscription, namely, genes transcribed in *Xenopus* oocytes (Jullien *et al*. 2014) and genes correlated with stem/progenitor cell transcript content in adult mouse organs (Kim *et al*. 2023). Enrichment analysis revealed that top 3000 highly transcribed genes from both sets are enriched among genes highly transcribed in the chicken oocyte (Figure 7 h). This finding suggests that different hypertranscription systems have similar regulatory mechanisms.

Next, to reveal general patterns of transcriptional regulation in oocytes at the lampbrush stage, all genes were ordered based on their transcript levels in chicken oocyte. Enrichment analysis was used to find key transcription regulators from ChEA 2022 database and histone modifications from ENCODE Histone Modifications 2015 database, which are associated with subsets of genes with the highest and the lowest levels in chicken oocyte. Clusters of tissue-specific genes from heart (o) and testis (p) were used to validate the results of the enrichment analysis.

Among the top 10 transcription factors associated with top 3000 genes highly transcribed in the chicken oocyte, there were the transcription factors RUNX2, known to be involved in early development, MYC factors, which promote global hypertranscription in the embryonic germ line (Percharde *et al*. 2017b), YY1, known to be involved in the formation of enhancer-promoter loops (Weintraub *et al*. 2017), and the H3K4-specific histone demethylase KDM5B (Supplementary Data Set 6). We noticed that a similar set of transcription regulators was associated with highly abundant transcripts in mouse adult stem/progenitor cells, which are characterised by a global transcriptome upregulation (Figure 7 i) (Kim *et al*. 2023).

Among the top 10 transcription regulators associated with 3000 genes with the lowest transcript level in chicken oocyte cytoplasm, there were components of polycomb repressive complexes PRC1 (RNF2, BMI1, CBX2, PHC1) and PRC2 (SUZ12, JARID2, EZH2, EED, MTF2) (Supplementary Data Set 6), which are known to be involved in chromatin compactisation and gene silencing via deposition of histone mark H3K27me3.

### Sex chromosome gene expression

We also checked the pattern of expression of 29 protein-coding genes retained on the chicken W chromosome (Bellott *et al*. 2017). According to RNA-seq data, most W chromosome genes are expressed at the lampbrush stage of oogenesis (Figure 2 b, d), presumably on the very small transcription loops at the borders of the compact chromomeres (Figure 2 c, c’). This pattern is consistent with the expression of a number of W chromosome genes in relation to female sex determination and adult reproductive system (Rallabandi *et al*. 2019). In contrast, certain genes involved in male gonadal differentiation, including the *DMRT1* gene, are not transcribed on the lampbrush chromosome Z in adult females (Figure 2 e).

In females, male hypermethylated repetitive region *MHM1* gives rise to long non-coding RNAs that play a role in the regulation of the *DMRT1* gene and gonadal sex differentiation (Melamed and Arnold 2007). It was previously shown that *MHM1* region close to the *DMRT1* gene locus is transcribed on a pair of loops on chicken ZW lampbrush bivalent (Teranishi *et al*. 2001). We observed polyadenylated nuclear retained transcript from the MHM1 locus that overlaps with the long non-coding RNA gene ENSGALG00000051419 (Figure 2 f). Consistent with this, the *MHM1* region is not hypermethylated in chicken lampbrush stage oocytes (Figure 2 g).

### Telomerase reverse transcriptase and telomerase RNA expression

Comparative analysis of transcriptome data allows reconstruction of the spatio-temporal pattern of telomerase component expression during chicken oogenesis and embryogenesis. Specifically, telomerase reverse transcriptase gene *TERT* is expressed in the chicken oocyte nucleus, with mRNA being found in the oocyte cytoplasm and then in the zygote and EGK.I-EGK.VIII embryos, thus representing maternal RNA (Figure S10 a). Telomerase RNA (TR) gene is found in a single locus between *MYNN* and *MECOM* genes on GGA9 and corresponds to the TR sequence (AY312571) described by (Delany and Daniels 2003). TR RNA is highly abundant in the chicken oocyte nucleus and is partially RNase R resistant (Figure S10 b). Interestingly, TERT and TR transcript levels are the highest in early differentiating embryos (O’Hare and Delany 2005). In chicken oocytes, expression of both components of telomerase correlates with the presence of telomeric non-coding RNA (TERRA RNA) in nuclear RNA samples (Figure 8 h, i). Previously, G-rich nascent telomere repeat transcripts were found at the ends of chicken lampbrush chromosomes by RNA-FISH (Solovei *et al*. 1994; Kulikova *et al*. 2016). We suggest that nascent telomere repeat transcripts may shield the telomere ends of lampbrush chromosomes from telomerase activity by pairing with the template region of telomerase RNA; however the telomerase activity would be required after fertilisation and early stages of embryogenesis.

**Figure 8.**
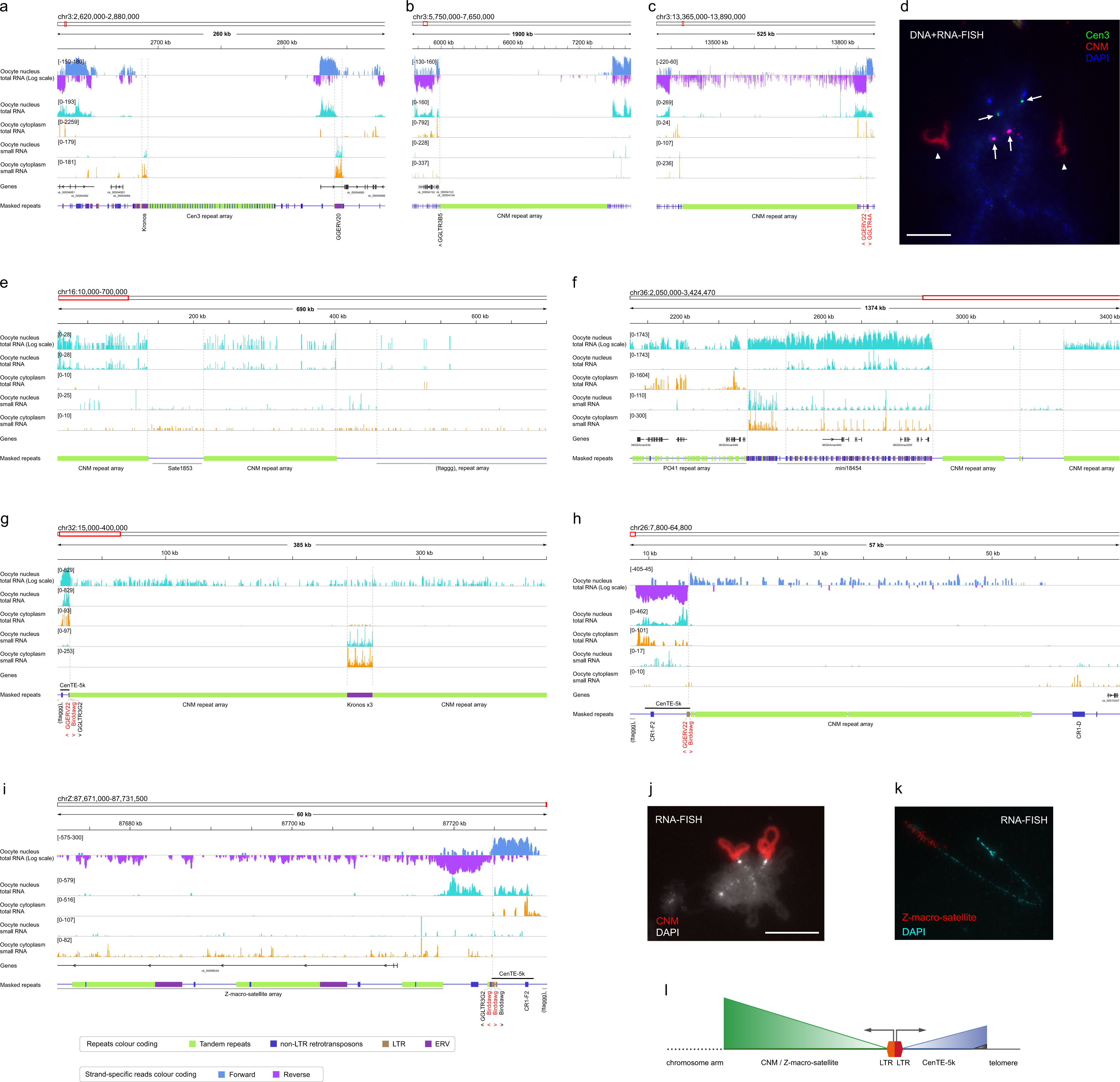
Chicken oocyte nuclear and cytoplasmic RNA profiles within centromere and subtelomere regions of chromosomes along the telomere-to-telomere chicken genome assembly. **a.** Overview of the centromere region of chicken chromosome 3. **b, c.** Overview of the regions containing non-centromeric arrays of CNM repeat on chicken chromosome 3. **d.** Visualisation of non-transcribed centromere and CNM repeat arrays as well as transcribed CNM repeat array on GGA3 using DNA+RNA-FISH with Cen3 probe and CNMneg oligonucleotide. Chromomeres in case of non-transcribed regions (arrows) and lateral loops in case of transcribed regions (arrowheads) are specifically labelled. **e-g.** Overview of the centromere regions of chicken chromosomes GGA16, GGA36 and GGA32. **h, i.** Overview of the subtelomere regions of chicken chromosomes GGA26 and GGAZ. Coverage tracks of total RNA and small RNA from the oocyte nucleus (cyan) and total RNA and small RNA from the oocyte cytoplasm (orange) of lampbrush stage oocytes visualised by IGV browser; count range limits are given in square brackets. For total RNA from the nucleus an additional coverage track is shown on a logarithmic scale; on panels a-c, h and i this additional coverage track visualizes strand-specific read counts. The Masked repeats track visualizes the positions of tandem repeats, non-LTR retrotransposons, LTRs and ERVs from the Repbase library of chicken repeats. The names of the selected repeats are shown below the track. The orientation of the selected LTR-containing elements is indicated by the symbols ‘>’ for forward and ‘<’ for reverse strand. Black horizontal bars indicate positions of the CenTe-5k sequence. Grey horizontal bars indicate positions of selected tandem repeat arrays. Vertical dotted lines indicate the position of the boundaries of selected transcription peaks. The names of LTR-containing elements whose position coincides with the boundaries of transcription peaks are shown in red. **j, k.** Visualisation of nascent transcripts of CNM and Z-macro-satellite tandem repeats. RNA-FISH with CNMneg oligonucleotide (**j**) or Z-macro-satellite specific probe (**k**) (red); lateral loops are specifically labelled. **l.** Organisation of transcription units at subtelomeric regions of chicken chromosomes showing that transcription of telomere-derived RNAs is initiated at adjacent LTR retrotransposons.

### Transcripts of tandem and interspersed repeats

To estimate the transcriptional activity at centromeres on all chicken chromosomes, including sex and dot chromosomes, during the diplotene stage of oogenesis, we mapped the oocyte total strand-specific RNA-seq data to the complete telomere-to-telomere assembly of the chicken genome. These data were compared with the previous RNA-FISH visualisation of nascent transcripts of tandem repeats on isolated chicken lampbrush chromosomes.

Mature chicken centromeres with chromosome-specific tandem repeats Cen1, Cen2, Cen3, Cen4, Cen7, Cen8 and Cen11, which interact with CENP-A (Shang *et al*. 2010), lie adjacent to the centromere granule in lampbrush chromosomes, occupying two distinct chromomeres (Krasikova *et al*. 2012). According to RNA FISH, the centromere clusters of these repeats are not transcribed on the lampbrush lateral loops (Krasikova *et al*. 2012). In full agreement with these data, complete silencing of the Cen1, Cen2, Cen3, Cen4, Cen7, Cen8 and Cen11 repeat arrays on GGA1-4, 7-8 and 11 was observed in oocyte nuclear and cytoplasmic RNA fractions (Figure 8 a, d). The results obtained are consistent with our model of centromere domain organisation, in which non-coding regulatory RNAs arise from peri-centromere regions while core centromeres are usually transcriptionally silent (Krasikova and Gaginskaia 2010).

Chicken chromosomes GGA5, GGAZ and GGA27 are characterised by recently evolved centromeres with unique centromere sequences (Shang *et al*. 2010). By aligning RNA-seq data for oocyte nuclear RNA fraction, we confirmed our previous RNA FISH data on the transcription of the non-tandemly-repetitive centromeres at the lampbrush stage of oogenesis (Krasikova *et al*. 2012) with most transcripts coming from certain genes.

The centromere regions of other chicken chromosomes, including dot chromosomes, have a more complex structure (Huang *et al*. 2023). The 41-bp repeats CNM and PO41 form higher-order repeats at the centromere regions of chicken acrocentric chromosomes (Huang *et al*. 2023). According to oocyte nuclear RNA-seq data, most CNM repeat clusters within centromeres of acrocentric chromosomes are transcriptionally silent (Figure 8 f). At the same time, the CNM arrays at the sub-telomeres, including those adjacent to the centromeres, are transcribed (Figure 8 e, f, j). The CNM repeat also has additional arrays on GGA3: the major CNM repeat array occupies a dense chromomere and the minor CNM repeat array forms a long transcriptional loop (Figure 8 d) (Krasikova *et al*. 2006). In complete agreement with RNA FISH mapping, in our nucleoplasmic RNA fraction the major CNM repeat array on GGA3 is silent (Figure 8 b), whereas the minor CNM repeat array is transcribed from one strand (Figure 8 c).

Tandem repeats embedded between CNM repeat arrays (Huang *et al*. 2023) often show high levels of transcripts according to oocyte nuclear RNA-seq data (e.g. in centromere regions of GGA25, GGA26, GGA28, GGA31, GGA36 chromosomes) (Figure 8 f). In many cases, RNA reads along simple tandem repeat arrays were only observed in nuclear RNA samples. In contrast, whole tissues RNA-seq profile did not show any transcripts along the centromere regions of chicken dot chromosomes (Huang *et al*. 2023).

The centromeric region of GGA32 dot chromosome is composed of several copies of the full-length Kronos retrotransposable element, surrounded by two CNM arrays (Huang *et al*. 2023). In our oocyte nuclear RNA-seq data, both Kronos and adjacent CNM repeat arrays on GGA32 undergo transcription during the lampbrush stage of oogenesis (Figure 8 g). In line with this, centromeric and pericentromeric RNAs have many functions at the centromeres and in the heterochromatin of the pericentromeres (Zhu *et al*. 2023).

W chromosome-specific *Xho*I-family repeat (Kodama *et al*. 1987), *Eco*RI-family repeat (Saitoh *et al*. 1991), and *Ssp*I-family repeat (Itoh and Mizuno 2002) do not demonstrate RNA reads in both nuclear and cytoplasmic oocyte RNA samples (Figure 2 a), which correlates with FISH mapping into chromomeres but not in the lateral loops (Solovei *et al*. 1998; Itoh and Mizuno 2002; Krasikova *et al*. 2006). Three major blocks of Z-Macro-Satellite on the long arm of chromosome Z are actively transcribed, with transcripts being found in oocyte nuclear RNA fractions (Figure 8 i, k). These results are consistent with previous data on the transcription of Z-Macro-Satellite on Z lampbrush chromosome, as revealed by FISH with nascent RNA on the lateral loops (Hori *et al*. 1996). In contrast, whole tissue RNA-seq profile did not show any transcripts along the hypermethylated region of the q arm of GGAZ (Huang *et al*. 2023).

The subtelomere regions of chicken chromosomes have similar genomic organisation. As shown by (Huang *et al*. 2023), almost half of the chromosomes in chicken genome share the CenTE-5k sequence, which separates (TTAGGG)n telomere and subtelomere repeat arrays. In our oocyte nuclear RNA samples, active transcription of subtelomere tandem repeat arrays was observed (Figure 8 h, i). Such transcription is initiated in opposite directions from long terminal repeats (LTRs) within the CenTE-5k sequence (Figure 8 l). For example, transcription initiation sites of subtelomere CNM repeat array at the left end of GGA26 and subtelomere Z-Macro-Satellite repeat at the 3’-end of the q arm of GGAZ coincide with Birddawg LTRs oriented towards the chromosome arm; initiation sites of transcription units towards telomere giving rise to telomeric repeat-containing RNA (TERRA) coincide with LTR containing element GGERV22 on GGA26 and Birddawg LTR on GGAZ, which are oriented towards the chromosome end (Figure 8 h, i). Thus, the data obtained are consistent with our previous suggestion that transcription of tandem repeats is initiated at the adjacent promoters of scattered LTR elements or even solo LTRs (LTR-activation hypothesis) (Deryusheva *et al*. 2007; Krasikova and Gaginskaia 2010; Trofimova and Krasikova 2016).

We then identified transcription start sites in the vicinity of the CNM arrays on chromosome The transcription start site located to the right of the minor CNM array coincides with the LTR containing element GGERV22 (Figure 8 c); transcription from this point continues throughout the minor CNM array. Adjacent to the GGERV22 element, is the GGLTR4A element, whose position coincides with the point at which transcription is directed away from the minor CNM array (Figure 8 c). Bidirectional transcription from LTR elements is typical for the transcription units in subtelomeric regions (Figure 8 g-i). It has previously been suggested that the region of GGA3 between the two CNM arrays is of microchromosomal origin (Zlotina *et al*. 2010). Thus, in this case, the transcribed short CNM repeat array corresponds to the subtelomeric region and the non-transcribed long CNM array corresponds to the centromeric region of this fused microchromosome.

### Chicken oocyte small RNAs reflect differential cytoplasmic and nuclear abundance

Four small RNA libraries from chicken lampbrush stage oocyte nucleus (two replicates) and cytoplasm (two replicates) were sequenced (Table S1) and classified according to known small non-coding chicken RNA species (Chan and Lowe 2009; Kozomara *et al*. 2019; Cunningham *et al*. 2022; Wang *et al*. 2022). Some copies of small housekeeping non-coding RNA genes are actively transcribed on lampbrush chromosomes, and their RNA products make up the majority of transcripts in the oocyte nucleus. Non-coding RNA species such as spliceosomal small nuclear RNAs (snRNAs), small nucleolar RNAs (snoRNAs), small Cajal body-specific RNAs (scaRNAs) were abundant in the oocyte nucleus but not cytoplasm, whereas tRNA and rRNA species were found in both oocyte compartments (Figure 9 b), consistent with known data on their intracellular localisation.

**Figure 9.**
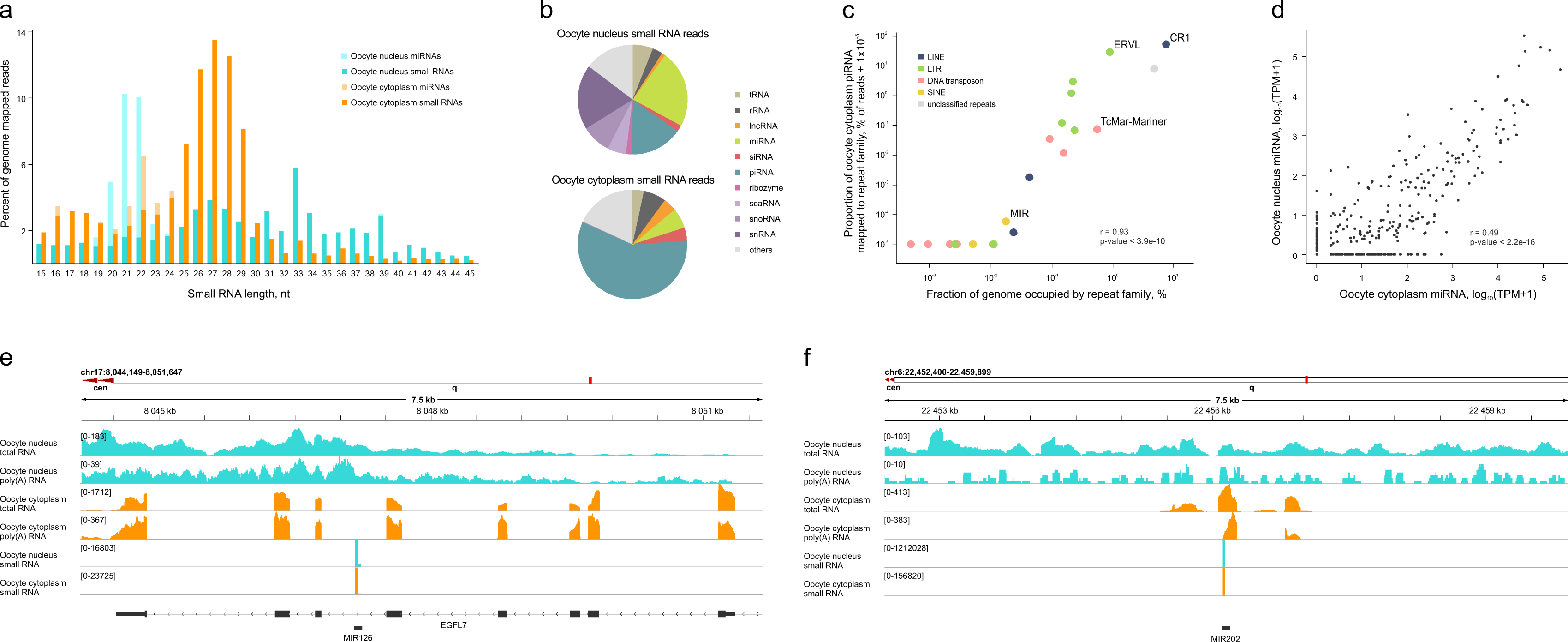
Chicken oocyte small RNA profile reflects differential cytoplasmic and nuclear abundance. **a.** Length distribution of reads in small RNA libraries from the chicken oocyte nucleus (cyan) and cytoplasm (orange); fractions of reads corresponding to known miRNAs are shown in lighter colours. **b.** Proportions of different classes of RNAs in small RNA libraries from the oocyte nuclei and cytoplasm. **c.** Scatter plot illustrating the relationship between the proportion of oocyte cytoplasmic piRNA reads mapped to different families of chicken interspersed repeats and the fraction of the chicken telomere-to-telomere genome assembly (GGswu1) occupied by corresponding family of interspersed repeats. Each dot represents a family of interspersed repeats; colours illustrate classes of interspersed repeats; both axes are shown in logarithmic scale. **d.** Scatter plot showing the correlation between miRNA levels in the oocyte nucleus and cytoplasm. **e, f.** Examples of transcribed host protein-coding and long non-coding RNA genes for expressed miRNAs. *EGFL7* host gene for embedded MIR126 and unannotated long non-coding RNA host gene for embedded MIR202 are shown. Coverage tracks of total RNA, poly(A) RNA and small RNA from the oocyte nuclei (cyan) and cytoplasm (orange) of lampbrush stage oocytes visualised by IGV browser along the gene body are shown; count range limits are given within square brackets. The exon-intron structure shown corresponds to one of the annotated transcript variants.

We found that the most abundant transcript in the oocyte nucleus is the 7SK RNA (Figure S11 d), which is known to play a key role in the negative control of transcription elongation (Studniarek *et al*. 2021). It is therefore possible that this small housekeeping RNA may have a role in shutting down transcription in the later stages of oocyte growth. In fact, in the late oocyte nucleus, the proportion of many small housekeeping RNAs in the total transcriptome increases compared to transcripts of protein-coding genes and lncRNAs.

In contrast to the absence of 18S and 28S rRNA in the oocyte nucleus, 5S rRNA is expressed within the lampbrush stage oocyte nucleus (Figure S11 b). Interestingly, most of the genes that encode the protein components of the ribosome subunits are also expressed by the oocyte itself.

U85 scaRNA is one of the most abundant RNAs in the oocyte nucleus and is not present in the oocyte cytoplasm. Chicken U85 scaRNA is transcribed with its host *NCAPD2* gene being processed from its intronic lariat. In addition, longer RNase R resistant stable intronic lariat bearing U85 scaRNA is observed in the nucleus (Figure S11 g). Another example of an RNase R resistant, stable intronic lariat bearing snoRNA in the oocyte nucleus is for the *HSPA8* intron-encoded SNORD14 (Figure S11 f), previously described in chicken DF1 cells (Talross *et al*. 2021). Such stable intronic lariats bearing snoRNA do not function as guide RNAs for rRNA and snRNA modification in contrast to mature snoRNA (Talross *et al*. 2021).

In contrast to U6 snRNA (Figure S11 a), U7 snRNA is not expressed in the oocyte (Figure S11 c), which correlates with the inactivation of the replication-dependent histone gene cluster on lampbrush chromosome 1 (Krasikova *et al*. 2012).

### Cytoplasmic short non-coding regulatory RNAs derived from nuclear transcripts

A total of 23% of the reads in the oocyte nucleus and 6% in the oocyte cytoplasm correspond to the mature microRNAs (miRNAs) (Figure 9 b). miRNA levels in biological replicates demonstrate a positive correlation (r = 0.97, p-value < 2.2e-16 for samples from both oocyte nucleus and oocyte cytoplasm) (Figure S12 a, b).

Out of 1164 miRNA genes annotated in the chicken genome (version GRCg6a), transcripts of 106 miRNAs were detected in the oocyte nucleus and 177 miRNAs were detected in the oocyte cytoplasm (Supplementary Data Set 7). Among them, 93 miRNAs present in both RNA fractions. Transcripts for both strands, −5p and −3p, were detected for 14 miRNAs in the oocyte nucleus and 54 miRNAs in the cytoplasm. miRNA levels in the nucleus and cytoplasm demonstrate a positive correlation (r = 0.49, p < 2.2e-16) (Figure 9 d), indicating transcription of miRNAs in the oocyte itself. Examples of transcribed host protein-coding and long non-coding RNA genes for expressed miRNA are shown on Figure 9 e-f.

To perform functional characterisation of miRNAs expressed in oocytes, 56 experimentally validated protein-encoding mRNA targets for miRNAs from the oocyte cytoplasm were found in the miRTarBase database (Supplementary Data Set 7i). Notably, 50 of these targets have transcripts in the oocyte itself, suggesting that the detected miRNAs can regulate the corresponding gene expression in the oocyte. For example, *RBFOX2* mRNA, known to be a direct target of miR-202 (Chen *et al*. 2016), is expressed in the oocyte nucleus, according to nuclear RNA-seq and RNA-FISH data (Figure S9 a). Enriched biological processes GO terms (P < 0.001) for miRNA targets are presented in Supplementary Data Set 7 and were mainly related to the regulation of transcription by RNA polymerase II, cell proliferation, cell differentiation, and organ development. A number of miRNAs, in particular miR-21, miR-125b and miR-202, shown here to be highly abundant in the oocytes (Figure 9 f), are known to be involved in the regulation of oogenesis and embryonic development (Wong *et al*. 2012; Alvi *et al*. 2021; Dehghan *et al*. 2021).

Post-transcriptional processing of the repeat transcripts may lead to the appearance of short regulatory RNAs such as endo-siRNA and piRNA. In our nuclear and cytoplasmic small RNA samples from chicken lampbrush stage oocytes, there are three major peaks in the length distribution of reads – at ∼22 nt, at ∼27 nt and at ∼33 nt (Figure 9 a). More than half of the peak at ∼22 nt corresponds to miRNA, as can be seen from the length distribution of reads after miRNA filtration (Figure 9 a). In human oocytes, in addition to miRNA and long piRNA, oocyte-specific piRNA (os-piRNA) of ∼20 nt in length appear due to presence of PIWIL3 Argonaute family protein (Yang *et al*. 2019). However, in mouse oocytes, instead of os-piRNA, endo-small interfering RNA of ∼22 nt in length appear due to absence of PIWIL3.

Similarly, *PIWIL3* gene was not found in chicken Entrez Gene and Uniprot databases. Therefore, we suggest that in the chicken oocyte small RNA length distribution after miRNA filtration, the rest of the small RNAs from the ∼22 nt peak corresponds to endo-siRNA, while the peak at ∼27 nt corresponds to piRNA. We also checked that *CIWI* (chicken *PIWIL1*) gene is expressed in chicken lampbrush stage oocytes.

Potential piRNAs from the ∼27 nt peak were the most abundant small RNAs in the cytoplasmic small RNA samples (Figure 9 b). A substantial proportion of the predicted piRNAs with the highest abundance in the oocyte cytoplasm correspond to the chicken piRNAs from the piRBase database (Wang *et al*. 2022) (Supplementary Data Set 8). We then analysed which repeats are targeted by predicted piRNAs in the chicken GGswu1 genome assembly (Table S4). The most abundant cytoplasmic and nuclear piRNAs target the CR1 (chicken LINE) retrotransposable element, the ERVL LTR element and the ERVK LTR element; small number of piRNAs target the TcMar-Mariner DNA transposon and almost no piRNAs target SINE elements (Figure 9 c).

## 3. Discussion

### Biological role of hypertranscription on the lateral loops of lampbrush chromosomes

Here we have carried out a systematic study of the spectrum of maternal RNAs in the chicken oocyte. We have established that at the lampbrush chromosome stage, chicken oocytes accumulate enormous amounts of mRNAs, long non-coding RNAs and small non-coding RNAs that are synthesised in the oocyte nucleus. Despite the giant size, chicken oocyte nucleus comprises only two chromosome sets. Thus to produce these vast number of RNAs extremely active transcription must happen at each of the >13000 expressed genes.

In the *Xenopus* oocyte, three strategies were thought to be realised to achieve this goal: the unusually high rate of nascent RNA synthesis, the months-long duration of transcription and the very high stability of the resulting mRNA (Gardner *et al*. 2012). Alternative way to produce large amounts of maternal transcripts is polyploidisation in the nurse cells surrounding the oocyte in animals with polygenomic type of oogenesis (Gall 2012). We conclude that in avian oocytes, hypertranscription on the lateral loops of giant lampbrush chromosomes is a primary mechanism to synthesise large amounts of maternal RNAs for each of the transcribed genes.

### Special features of hypertranscription during the lampbrush stage of oogenesis

Genome expression can be regulated by the speed of transcription (Muniz *et al*. 2021). The rate of nascent RNA synthesis is unusually high in lampbrush chromosomes (Gall and Callan 1962; Izawa *et al*. 1963; Callebaut 1973; Hartley and Callan 1978; Deryusheva *et al*. 2007; Kulikova *et al*. 2016; Morgan 2018). This is most likely achieved by the tight coverage of transcribed genes by elongating RNA polymerases as shown by chromosome spreading according to the Miller technique (Scheer *et al*. 1976). An increase in the rate of elongation by RNA polymerase II also cannot be ruled out. The rate of transcript production can be also adjustable depending on the particular gene. At the same time, studies on incorporation of RNA precursors into the nascent transcripts in lampbrush chromosomes of amphibians and birds demonstrated uniform coverage of simple lateral loops; the only exceptions were a few rare complex loops with unusual morphology of the RNP matrix, which showed a lower rate of incorporation of RNA precursors (Gall and Callan 1962). More detailed studies are needed to establish a relationship between the level of gene expression, calculated from intron read counts, the density of RNA polymerases along transcription units and the extension of transcription loops in lampbrush chromosomes.

The speed of RNA polymerase can be also measured by a gradient of reads from the 5′ to the 3′ end of the intron (Jonkers *et al*. 2014; Debes *et al*. 2023). Accordingly, we observed a saw-tooth pattern of reads along the transcribed genes, which was particularly evident for very long genes with lengthy introns.

RNA-polymerase speed can influence the transcriptome profile (Muniz *et al*. 2021). We have shown that transcription on the majority of normal lateral loops on avian lampbrush chromosomes is initiated at the promoters of active genes; transcription start sites often correspond to CpG islands. This is consistent with previous findings on hypomethylation of the majority of promoters and CpG islands in single-cell methylome data of the chicken diplotene oocytes (Nurislamov *et al*. 2022). Transcript boundaries coincide with gene boundaries and the transcribed strand corresponds to the orientation of a gene.

Polyadenylation of the transcripts means that RNA polymerase II successfully proceeds through the entire length of the transcription unit. The appearance of correctly spliced mRNAs and long non-coding RNAs in the cytoplasm for the transcribed genes indicates on normal splicing. However, it would be interesting to investigate the effect of the high rate of RNA synthesis in lampbrush chromosomes on the alternative splicing and alternative sites of RNA polyadenylation.

Transcriptional read-through sometimes accompanies increase in RNA polymerase speed affecting expression of downstream genes (Muniz *et al*. 2017) and remodelling genome architecture (Heinz *et al*. 2018). Read-through transcription, which is initiated at histone gene promoters and continues through downstream histone genes or even satellite DNA, has previously been demonstrated in *Notophthalmus* and *Xenopus* oocytes (Diaz and Gall 1985; Masi and Johnson 2003). We observed only rare examples of read-through transcription on chicken lampbrush chromosomes, with most gene transcripts being normally polyadenylated. Furthermore, with very few exceptions, individual loci are transcribed from only one strand. We propose that, due to hypertranscription, gaps between transcriptional bursts, which potentially result in transcripts from overlapping genes, are very short and infrequent and therefore do not allow transcription of the expressed gene to be interrupted.

Higher RNA polymerase II velocity also directly correlates with the production of newly synthesised circular RNA (Zhang *et al*. 2016; Ragan *et al*. 2019). Comparison of the chicken oocyte nucleus and cytoplasm RNA profiles clearly showed the nuclear retention of certain introns in the form of sisRNAs resistant to RNase R treatment. Interestingly, ring-like structures associated with the nascent RNP fibrils were observed during electron microscopy of spread lampbrush chromosome preparations (Figure 14 in Scheer *et al*. 1976); these may correspond to sisRNAs being processed co-transcriptionally. sisRNA accumulation was previously demonstrated for *Xenopus* lampbrush stage oocyte nuclei (Gardner *et al*. 2012). Further studies of sisRNAs showed that they are usually in a circular form, and 9000 sisRNAs were found in the oocyte cytoplasm (Talhouarne and Gall 2014). sisRNAs have subsequently been identified in the cytoplasm of the somatic cells in a variety of vertebrate species (Talhouarne and Gall 2018).

The high number of certain intronic sequences stored within the oocyte nucleus is due to months of expression of corresponding genes on the lateral loops of lampbrush chromosomes as well as due to their unusual stability. Notably, the production of sisRNA generally does not interfere with host gene expression and the appearance of the corresponding mRNA. sisRNAs apparently play a role in the regulation of their host gene expression (Osman *et al*. 2016; Chan and Pek 2019). For example, a circular sisRNA ci-ankrd52 colocalises with the transcription sites of the host gene and positively regulates its transcription via association with elongating form of RNA polymerase II (Zhang *et al*. 2013). We propose that nuclear retained sisRNAs produced by a number of transcribed genes on the lateral loops of avian and amphibian lampbrush chromosomes via splicing-dependent mechanism can maintain the continued transcription of their host genes either at the lampbrush stage of oogenesis or during embryogenesis.

Elevated transcription in lampbrush chromosomes may also affect the 3D organisation of chromatin domains, which we will discuss elsewhere.

### Maternal RNAs synthesised on the lateral loops of lampbrush chromosomes

Strong overlap of mRNA and long non-coding RNA profiles in the oocyte nucleus and cytoplasm indicates transcription within the oocyte nucleus at the lampbrush stage. In particular, exons predominate in the oocyte cytoplasm, whereas introns of the same genes predominate in the oocyte nucleus. More than 95% of all reads from the oocyte cytoplasm were mapped within annotated genes. So in the oocyte, not everything is transcribed, but certain genes. Transcripts for ∼60% of protein-coding genes, ∼40% of lncRNAs and ∼15% of miRNAs were detected. The total number of genes transcribed in the oocyte itself is therefore comparable to that in other tissues.

The overlap between the oocyte and embryo RNA profiles before the major wave of zygotic genome activation indicates that maternal RNA synthesised by the oocyte itself and accumulated in its cytoplasm is inherited by the embryo. We assume that maternally deposited transcripts derived from the oocyte nucleus are required for the processes associated with oogenesis and early embryogenesis.

Among genes expressed in the oocyte nucleus, those with a broader expression profile predominate compared to genes whose transcripts were absent in the oocyte. Clusters of genes whose high transcript levels correlate with each other in the oocyte nucleus and cytoplasm are mainly associated with essential cellular processes relevant to the oocyte and embryo physiology. These processes include transcription, mRNA catabolism, piRNA metabolism, DNA repair, RNA splicing, protein synthesis, DNA replication, cell cycle progression and mitochondria functions. Notably, the gene cluster with a maximum of transcript levels in the oocyte cytoplasm is associated with the process of transcription, while the gene cluster with a maximum of expression in early embryogenesis is associated with the processes of mRNA catabolism, DNA replication and cell cycle progression.

We have also shown for the first time, that regulatory long non-coding RNAs are synthesised inside the oocyte nucleus during the lampbrush stage of oogenesis. Previously, lower levels of long non-coding RNAs compared to protein-coding gene transcripts have been reported for various organs (Jehl *et al*. 2020), whereas here we found higher levels of lncRNAs in the oocyte nucleus. Genes for lncRNAs were enriched in two clusters - the cluster of nuclear retained RNAs, suggesting their function in intranuclear events, and the cluster of genes not transcribed in the oocyte, consistent with their role as tissue-specific regulators activated at later stages of development (Sarropoulos *et al*. 2019).

In addition to the high rate of on-going transcription on the lateral loops of the lampbrush chromosomes inside the avian oocyte nucleus, a limited set of RNA species seem to be synthesised at the earlier stages of oogenesis. Among such RNA species are nulceolar rRNA and histone mRNAs that are highly abundant in the oocyte cytoplasm but not the nucleus. Indeed, early diplotene stage oocytes from adult hens poses active nucleolus that disintegrates at the beginning of the lampbrush stage (Davidian *et al*. 2023). At the same time, granulosa cells could be involved in supplying the oocyte with extraembryonic RNA under the perivitelline layer outside the germinal disc that is not used for the embryo development (Olszanska and Stepinska 2008).

### Regulation of hypertranscription of maternal RNA genes on lampbrush chromosomes

Our data suggests that transcription of the expressed set of genes at the lampbrush stage of oogenesis is positively controlled by KDM5B demethylase and the associated histone modification H3K4me3. KDM5B is involved in focusing H3K4 methylation marks near promoters and enhancers of active genes in embryonic stem cells including core pluripotency regulators (Kidder *et al*. 2014), activates transcriptional elongation along self-renewal-associated genes (Xie *et al*. 2011) and regulates nucleosome positioning and RNA polymerase II occupancy (Kurup *et al*. 2019). In chicken lampbrush chromosomes, H3K4me3 showed a punctate pattern at the bases of laterally extending loops (Krasikova *et al*. 2009).

At the same time, repression of the inactived genes in avian lampbrush chromosomes is controlled by epigenetic mechanisms involving Polycomb complexes and histone modification H3K27me3. This suggestion is supported by the accumulation of H3K27me3 in lampbrush chromomeres, with particular abundance in the regions of constitutive heterochromatin (Krasikova *et al*. 2009).

### Visualisation of nascent RNA synthesis and processing in monogenic and multigenic transcription loops

In addition, for a number of genes we directly visualised nascent transcripts on the lateral loops of the lampbrush chromosomes by RNA-FISH. Moreover, we argue that nuclear RNA-seq profile allows to predict the chromomere-loop pattern of lampbrush chromatin domains. Genomic regions containing a number of actively transcribed genes would form a transcription loop, while genomic regions without transcription units would be packed into chromomeres or chromatin knots. We described transcription loops for *RBFOX1*, *RBMS3*, *CCSER1*, *GRID2*, *INPP5A* and *PARD3B* genes, some of them being involved in embryo development and maternal RNA stability. Visualisation of co-transcriptional splicing in case of *RBFOX1* and *RBMS3* genes by light microscopy is accompanied by detection of correctly spliced maternal mRNAs deposited in the oocyte cytoplasm. In the late oocyte nucleus, transcriptional output tends to decrease while the boundaries of transcription units are maintained. This is consistent with RNA-FISH data showing the presence of nascent gene transcripts on the lateral loops, which shorten dramatically in late stage oocytes.

New examples of genes whose newly synthesised transcripts were detected in the RNP matrix of lateral loops of lampbrush chromosomes support previous findings. Nascent transcripts of the genes for the nucleoplasmin, cytokeratin and basal fibroblast growth factor were identified on the lateral loops of the *X. laevis* lampbrush chromosomes (Weber *et al*. 1989; Sallacz and Jantsch 2005). Recently, RNA-FISH with BAC clone based probes selected according to oocyte RNA-seq data identified transcripts overlapping with 13 protein-coding genes on axolotl (*Ambistoma mexicanum*) lampbrush chromosomes (Keinath *et al*. 2021). By RNA-FISH with BAC clones we also revealed transcripts from 13 protein-coding genes and 2 non-coding RNA genes on the lateral loops of chicken lampbrush chromosomes (Krasikova *et al*. 2012; Kulikova *et al*. 2022). For these genes, we also confirmed the presence of transcripts in the oocyte nuclear and cytoplasmic RNA fractions (Table S3).

Therefore, lampbrush chromosomes provide an especially useful system for studying thousands of monogenic and multigenic transcription loops as well as the packaging of transcriptionally active chromatin and nascent RNAs. Transcription loops showing the progression of nascent RNA synthesis and splicing have been described for individual hyperactively expressed long genes in interphase nuclei of somatic cells (Leidescher *et al*. 2022; Ullrich *et al*. 2023). Extension of the transcription loops in interphase nuclei is directly dependent on the expression level of the gene and most likely reflects the frequency and duration of transcriptional bursts demonstrating high similarity to lampbrush lateral loops.

### Short regulatory piRNAs can participate in inactivation of interspersed repeats during early embryogenesis

Lampbrush chromosomes can transcribe not only protein-coding and long non-coding RNA genes, but also miRNAs, both as part of the host gene transcript and as independent genes. Here we characterise for the first time a full set of small non-coding RNAs from the cytoplasm and nucleus of lampbrush stage oocytes, including miRNAs and piRNAs. Several studies have characterised miRNA profiles in ovarian tissue from chickens of different ages, including sexually mature females (Kang *et al*. 2013). miRNA profile was also characterised in chicken ovarian follicles including prehierarchical white follicles (Oclon and Hrabia 2021). However, in addition to the oocyte, other cell types were present in these tissues including granulosa cells, theca cells and blood vessels. miRNAs identified in the chicken lampbrush stage oocyte cytoplasm in our study are involved into the regulation of transcription by RNA polymerase II, cell proliferation, cell differentiation, and organ development and could be also utilised during embryogenesis.

About 254000 piRNAs in chicken enucleated oocytes and their potential targets were characterised in this study. The most abundant piRNA species that we found in chicken oocyte target CR1 and LTR containing retrotransposable elements. CR1 elements are the most numerous repeats in the chicken genome; CR1 represents autonomous LINE retrotransposon with few functional “mother” elements (Wicker *et al*. 2005). Several full-length LTR containing elements also remain intact in the chicken genome and possess potentially functional ORFs (Wicker *et al*. 2005). Thus CR1 and LTR containing elements should be targeted by piRNAs to prevent their spreading (Sun *et al*. 2023). Among autonomous DNA transposons, no functional *Mariner* or Charlie elements encoding a transposase have been characterised in the chicken genome (Wicker *et al*. 2005; Kordis 2009). Accordingly, in the growing chicken oocyte, we found only a small number of unique piRNA species that target these DNA transposons.

The oocyte cytoplasm was highly enriched in piRNAs, indicating their accumulation; this set of piRNAs may be involved in the inactivation of retrotransposable elements during embryogenesis (Krasikova and Fedorov 2016). Transposon-derived piRNA-like small RNAs (pilRNA) targeting preferentially LTR and CR1 elements were characterised in EGK.X chicken embryos (Shao *et al*. 2012). However, at this stage of embryogenesis, pilRNA were of post-zygotic origin and localised specifically to the primordial germ cells. Further studies are needed to analyse the small RNA profile in early chicken embryos to identify maternally inherited miRNAs and piRNAs.

We also consider transcripts from tandem repeats as long non-coding RNA. Here, we show that in chicken lampbrush chromosomes transcription of telomere-derived RNAs, including telomeric repeat-containing RNA (TERRA) and subtelomere repeat-containing RNA, is initiated at neighbouring LTR elements, suggesting a role for active retrotransposons in the regulation of telomere non-coding RNA synthesis. Such genomic organisation of subtelomere regions of chicken chromosomes corresponds to the arrangement of transcription loops at the ends of lampbrush chromosomes (Solovei *et al*. 1994; Hori *et al*. 1996). As shown by RNA-FISH, some CNM repeat arrays are transcribed from one strand, while the others are transcribed from another strand (Krasikova *et al*. 2006; Deryusheva *et al*. 2007). We propose that transcription of subtelomere repeats (CNM, PO41 or Z-macro-satellite in chicken) in the opposite direction to the telomere protects the telomere from read-through transcription of genes located on the chromosome arms. We also suggest that tandem repeat transcripts in chicken lampbrush-stage oocytes are utilised to produce short regulatory siRNAs that can participate in heterochromatin formation during early embryogenesis (Krasikova and Fedorov 2016).

Therefore, our chicken oocyte nucleus and cytoplasm transcriptome analysis provides rigorous proof on the role of lampbrush chromosomes in the maternal RNA synthesis. Comparison of the RNAs stored in the oocyte with early embryonic RNAs suggests that mRNAs and long non-coding RNAs synthesised on the lateral loops of lampbrush chromosomes inside the oocyte nucleus play a role in both oocyte maturation and embryo development, as well as in regulating embryonic gene expression.

Transcription that occurs at the lampbrush stage of oogenesis is generally similar to that of many other cell types in terms of basic regulatory mechanisms and the spectrum of sequences that are transcribed. However due to the unusually high intensity of transcription and giant size, lampbrush chromosomes can be used as a high-resolution model to study gene expression, regulation of transcription, RNA processing, transcription loop dynamics and chromatin domain formation. Given that hypertranscription is typical not only of the oocyte nucleus, but also of embryonic germline and adult stem cells (Percharde *et al*. 2017a), lampbrush chromosomes provide a valuable model for studying mechanisms of global transcriptome up-regulation.

## Supporting information

Supplementary Data Sets

## Acknowledgments

This manuscript is dedicated to the memory of Herbert Macgregor. The research was supported by the Russian Science Foundation (grant #19-74-20075) and was performed using the equipment of the Genomics Core Facility (Skoltech) and Resource Center “Molecular and Cell Technologies” (Saint-Petersburg State University). We are grateful to Alexei Popov (Genomics Core Facility, Skoltech) for initial bioinformatic analysis of the RNA-seq data.

## Author contributions

AK - conceptualisation, study design, supervision of the project and funding acquisition; TK, AM, Ant F - RNA isolation; NM, Ann F - RNA-seq workflow; MS - bioinformatic analysis of RNA-seq data; Ant F - transcriptomic data visualisation and enrichment analysis; AK, TK, Ant F - visual RNA-seq data analysis; TK, VB - FISH on lampbrush chromosomes; TK, VB - microscopic experiments; TK, AK - image analysis; AK, Ant F - wrote the manuscript; TK, MS - revised the manuscript; all co-authors approved the final version of the manuscript.

## Competing interests

The authors declare no competing interests

## Data availability

RNA-seq data is available at the NCBI Sequence Read Archive (SRA) via accession numbers provided at the Supplementary Table S1. Any additional data in support of the findings of this study can be obtained from the corresponding author upon reasonable request.

## Supplementary Materials

### Supplementary Figures

**Figure S1.** Total RNA profiles from chicken lampbrush-stage oocyte nucleus and oocyte cytoplasm against the telomere-to-telomere chicken chromosome assembly.

**Figure S2.** The fraction of the chicken telomere-to-telomere genome assembly (GGswu1) covered at a given depth by RNA-seq reads fromRNA libraries of lampbrush stage oocyte nuclei and cytoplasm.

**Figure S3.** Nuclear and cytoplasmic total and poly(A) enriched RNA sequences from protein-coding and non-coding RNA genes.

**Figure S4.** Gradient in total RNA read coverage from 5’ to 3’ end of the introns of the DMD, LSAMP, BEND5, and AUTS2 genes, reflecting elongation by RNA polymerase II and co-transcriptional splicing.

**Figure S5.** Most genomic regions in chicken lampbrush stage oocytes are transcribed from one strand.

**Figure S6.** Correlation between poly(A) RNA and total RNA reads from (a) oocyte nucleus and (b) oocyte cytoplasm.

**Figure S7.** Visualisation of nascent pre-mRNA from individual transcribed protein-coding genes (CSMD3, INPPA5 and PARD3B) by RNA-FISH on lampbrush chromosomes.

**Figure S8.** At the post-lampbrush chromosome stage, the transcription loops with nascent gene transcripts become substantially shorter.

**Figure S9.** Comparison of RNA-seq data and RNA-FISH pattern along the genomic regions containing multiple transcribed genes and forming lateral loops.

**Figure S10.** Telomerase reverse transcriptase and telomerase RNA expression during oogenesis and early embryogenesis.

**Figure S11.** Examples of transcribing and non-transcribing small housekeeping RNA genes.

**Figure S12.** Correlation between miRNA (a, b) and piRNA (c, d) reads in biological replicates from oocyte nucleus (a, c) and cytoplasm (b, d).

### Supplementary Tables

**Table S1.** RNA-seq data available from the NCBI Sequence Read Archive (SRA) using the accession numbers provided.

**Table S2.** The list of BAC clones containing fragments of chicken genomic DNA from the CHORI-261 library that were used as DNA-probes for FISH.

**Table S3.** The list of annotated gene sequences (according to galGal6 chicken genome assembly) whose transcriptional status was verified by both RNA-FISH on lampbrush chromosomes and RNA-seq of oocyte nuclear and cytoplasmic RNA.

**Table S4.** Targets of predicted piRNAs in the chicken oocyte nucleus and oocyte cytoplasm.

### Supplementary Data Sets

**Supplementary Data Set 1.** RNA-seq data of levels of gene transcripts in total RNA samples from chicken oocyte nucleus and oocyte cytoplasm.

**Supplementary Data Set 2.** List of gene regions for which transcripts from both strands were revealed. Contains information about the coordinates of overlapping transcripts, the length of the overlap, and the genes intersected by the overlapping transcripts. Minimum length of overlap is 100 nucleotides.

**Supplementary Data Set 3.** Lists of predicted circRNAs in different RNA samples. Includes information on the coordinates of predicted circRNAs, their length, number of reads spanning the back-splice junction and gene regions overlapped by circRNA transcripts.

i. Total RNA from oocyte nucleus treated with RNAse R.
ii. Total RNA from oocyte nucleus.
iii. Poly(A) RNA from oocyte nucleus.

**Supplementary Data Set 4.** Enriched biological processes GO terms associated with clusters of genes co-expressed in chicken oocyte and early embryo.

**Supplementary Data Set 5.** Enriched biological processes GO terms associated with clusters of genes co-expressed in chicken oocyte and adult organs.

**Supplementary Data Set 6.** Enriched transcription factors (i) and histone modifications (ii) associated with subsets of genes with the highest and the lowest transcript levels in chicken oocyte. Clusters of tissue-specific genes from heart (o) and testis (p), were used to control results of enrichment analysis. Expression dataset for adult chicken organs was obtained from literature (Bush et al., 2018).

**Supplementary Data Set 7.** Data on miRNA abundance in the chicken oocyte nucleus and cytoplasm.

i. RNA-seq data of miRNA levels in small RNA samples from chicken oocyte nucleus and oocyte cytoplasm.
ii. Experimentally validated protein-coding mRNA targets of oocyte cytoplasmic miRNAs retrieved from miRTarBase (https://mirtarbase.cuhk.edu.cn/).
iii. Enriched biological processes GO terms associated with experimentally validated protein-encoding mRNA targets for miRNAs.

**Supplementary Data Set 8.** Data on abundance of predicted piRNAs in the chicken oocyte nucleus and cytoplasm.

## Notes

### Competing Interest Statement

The authors have declared no competing interest.

