## Supplementary Materials for "The first chicken oocyte nucleus whole transcriptomic profile defines the spectrum of maternal mRNA and non-coding RNA genes transcribed by the lampbrush chromosomes"

Table S4. Targets of predicted piRNAs in the chicken oocyte nucleus and oocyte cytoplasm.

***List of References for the Supplementary Materials***

***Supplementary Data Sets***

List of Supplementary Data Sets.

### Supplementary figures

**Figure S1. Total RNA profiles from chicken lampbrush-stage oocyte nucleus and oocyte cytoplasm against the telomere-to-telomere chicken chromosome assembly.** W chromosome total RNA profile is shown on the Figure 2 a. Total RNA profiles for the EGK.III and EGK.X stages of embryogenesis along the complete chicken chromosome assemblies are also shown. RNA-seq data for EGK.III and EGK.X stages of embryogenesis was taken from (Hwang *et al.* 2018a).

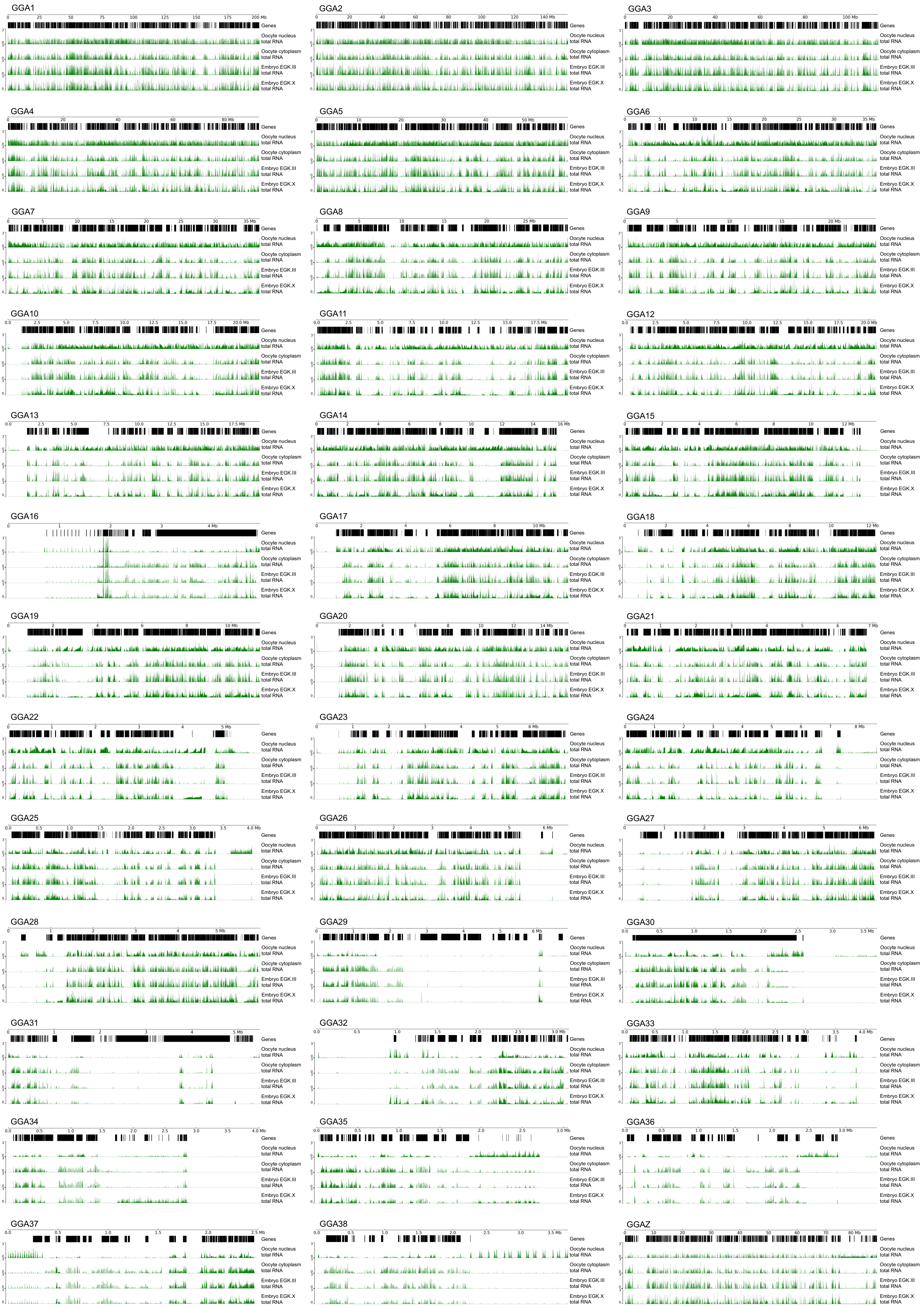
GGA1
GGA2
GGA3
GGA4
GGA5
GGA6
GGA7
GGA8
GGA9
GGA10
GGA11
GGA12
GGA13
GGA14
GGA15
GGA16
GGA17
GGA18
GGA19
GGA20
GGA21
GGA22
GGA23
GGA24
GGA25
GGA26
GGA27
GGA28
GGA29
GGA30
GGA31
GGA32
GGA33
GGA34
GGA35
GGA36
GGA37
GGA38
GGAZ
Genes
Oocyte nucleus total RNA
Oocyte cytoplasm total RNA
Embryo EGK.III total RNA
Embryo EGK.X total RNA

**Figure S2. The fraction of the chicken telomere-to-telomere genome assembly (GGswu1) covered at a given depth by RNA-seq reads from RNA libraries of lampbrush stage oocyte nuclei and cytoplasm.**

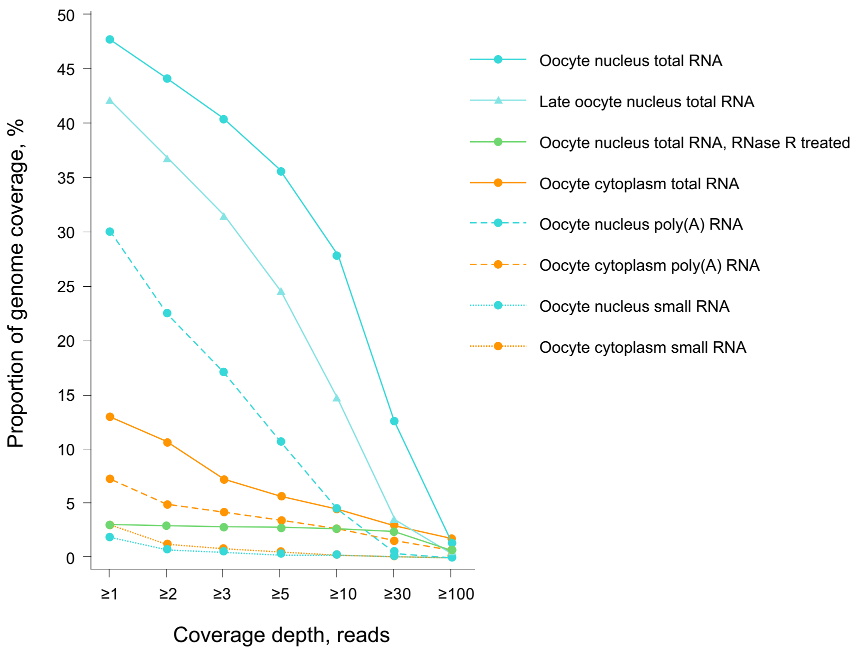

Proportion of genome coverage, %
50
45
40
35
30
25
20
15
10
5
0
≥1
≥2
≥3
≥5
≥10
≥30
≥100
Coverage depth, reads
Oocyte nucleus total RNA
Late oocyte nucleus total RNA
Oocyte nucleus total RNA, RNase R treated
Oocyte cytoplasm total RNA
Oocyte nucleus poly(A) RNA
Oocyte cytoplasm poly(A) RNA
Oocyte nucleus small RNA
Oocyte cytoplasm small RNA

**Figure S3. Nuclear and cytoplasmic total and poly(A) enriched RNA sequences from protein-coding and non-coding RNA genes. a-f.** Oocyte cytoplasmic mRNA from *PTPA*, *NOTCH1*, *PTBP1, MYC,* and *SOX3* genes and long non-coding RNA from ENSGALG00000048648 gene consist primarily of spliced exons. Oocyte nuclear RNA contains nascent RNA and products of RNA processing for the same genes. **g, h. Stable intronic sequence RNAs (sisRNAs) resistant to RNase R treatment spliced from the nascent pre-mRNA transcript.** Examples of sisRNA-producing genes: *NPHP4* and *XPO7*. Coverage tracks of total and poly(A) RNA from the oocyte nuclei (cyan) and cytoplasm (orange) of lampbrush stage oocytes visualised by IGV browser along the gene body are shown; count range limits are given within square brackets. Coverage track of total RNA from the oocyte nuclei after RNase R treatment reveals sisRNAs (green). The exon-intron structure shown corresponds to one of the annotated transcript variants.

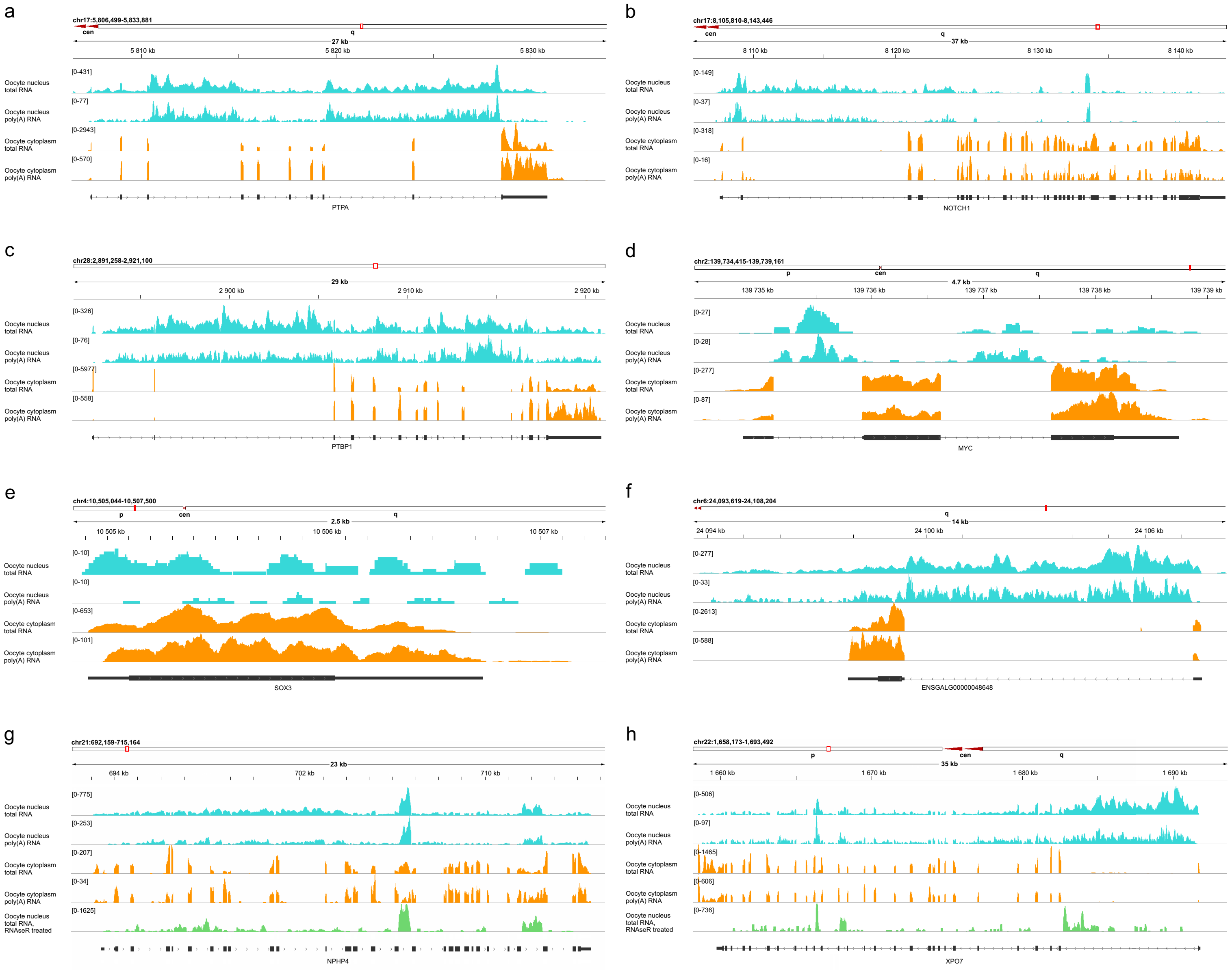
a
chr17:5,806,499-5,833,881
cen
q
27 kb
5 810 kb
5 820 kb
5 830 kb
Oocyte nucleus total RNA
[0-431]
Oocyte nucleus poly(A) RNA
[0-77]
Oocyte cytoplasm total RNA
[0-2943]
Oocyte cytoplasm poly(A) RNA
[0-570]
PTPA
b
chr17:8,105,810-8,143,446
cen
q
37 kb
8 110 kb
8 120 kb
8 130 kb
8 140 kb
[0-149]
[0-37]
[0-318]
[0-16]
NOTCH1
c
chr28:2,891,258-2,921,100
29 kb
2 900 kb
2 910 kb
2 920 kb
[0-326]
[0-76]
[0-5977]
[0-558]
PTBP1
d
chr2:139,734,415-139,739,161
p
cen
q
4.7 kb
139 735 kb
139 736 kb
139 737 kb
139 738 kb
139 739 kb
[0-27]
[0-28]
[0-277]
[0-87]
MYC
e
chr4:10,505,044-10,507,500
p
cen
q
2.5 kb
10 505 kb
10 506 kb
10 507 kb
[0-10]
[0-10]
[0-653]
[0-101]
SOX3
f
chr6:24,093,619-24,108,204
q
14 kb
24 094 kb
24 100 kb
24 106 kb
[0-277]
[0-33]
[0-2613]
[0-588]
ENSGALG00000048648
g
chr21:692,159-715,164
23 kb
694 kb
702 kb
710 kb
[0-775]
[0-253]
[0-207]
[0-34]
Oocyte nucleus total RNA, RNAseR treated
[0-1625]
NPHP4
h
chr22:1,658,173-1,693,492
p
cen
q
35 kb
1 660 kb
1 670 kb
1 680 kb
1 690 kb
[0-506]
[0-97]
[0-1465]
[0-606]
[0-736]
XPO7

**Figure S4. Gradient in total RNA read coverage from 5' to 3' end of the introns of the *DMD*, *LSAMP*, *BEND5*, and *AUTS2* genes, reflecting elongation by RNA polymerase II and co-transcriptional splicing (a-d).** The sawtooth pattern, reflecting gradients in RNA density along introns, is more evident in the nuclear RNA fraction from oocytes at the lampbrush chromosome stage (cyan) and less evident in the nuclear RNA fraction from oocytes at the post-lampbrush chromosome stage (late oocyte nucleus) (light cyan). Oocyte cytoplasm (orange) RNA-seq profile is also shown. The exon-intron structure shown corresponds to one of the annotated transcript variants.

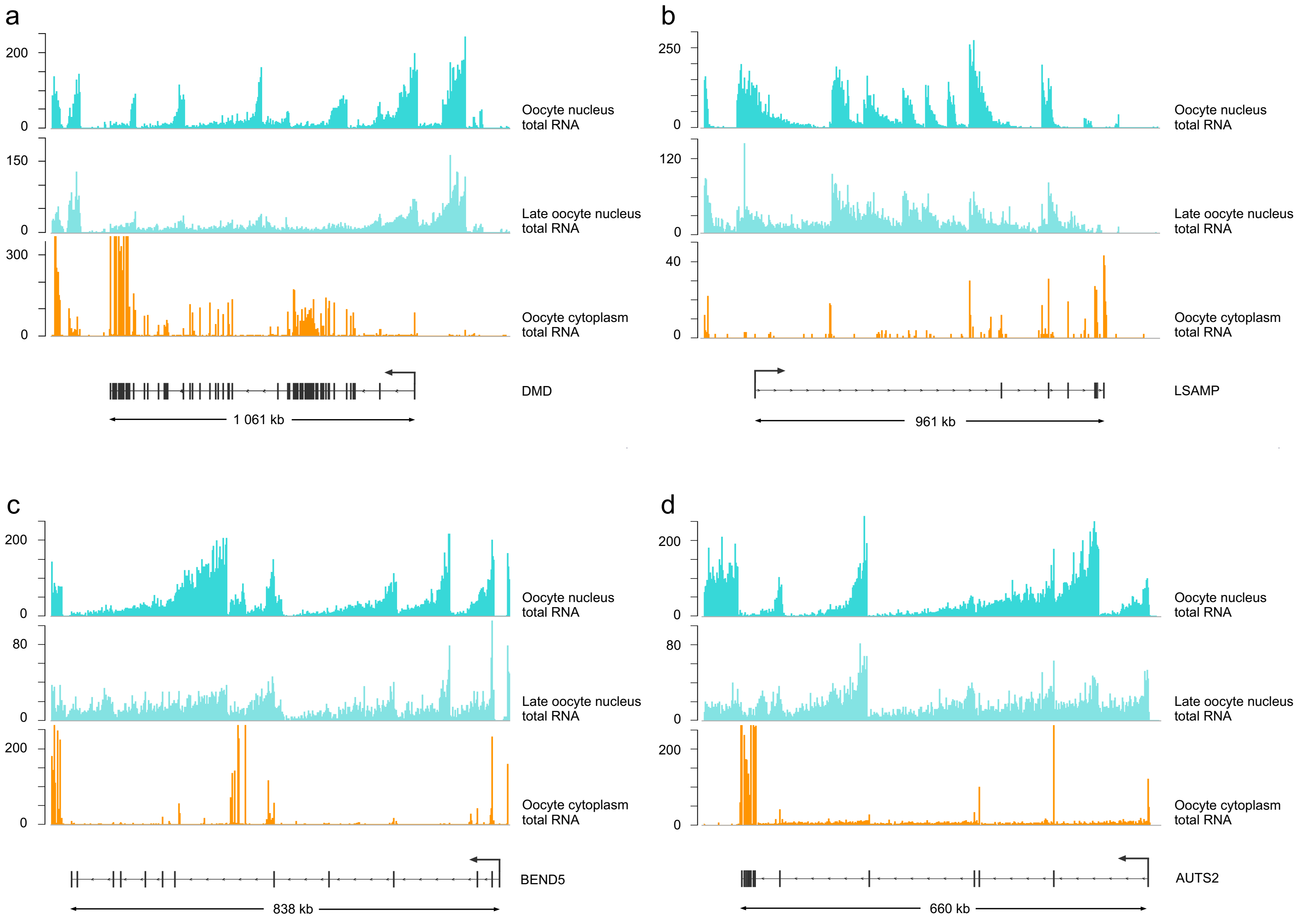
a
200
0
Oocyte nucleus
total RNA
150
0
Late oocyte nucleus
total RNA
300
0
Oocyte cytoplasm
total RNA
DMD
1 061 kb
b
250
0
Oocyte nucleus
total RNA
120
0
Late oocyte nucleus
total RNA
40
0
Oocyte cytoplasm
total RNA
LSAMP
961 kb
c
200
0
Oocyte nucleus
total RNA
80
0
Late oocyte nucleus
total RNA
200
0
Oocyte cytoplasm
total RNA
BEND5
838 kb
d
200
0
Oocyte nucleus
total RNA
80
0
Late oocyte nucleus
total RNA
200
0
Oocyte cytoplasm
total RNA
AUTS2
660 kb

**Figure S5. Most genomic regions in chicken lampbrush stage oocytes are transcribed from one strand.**

**a-c.** Examples of genomic regions with divergent and convergent gene pairs transcribed from one strand: *USP5* and *CDCA3* genes (**a**), *PHC1* and *M6PR* genes (**b**), *NEFL* and *NEFM* genes (**c**). Oocyte nuclear and cytoplasmic stranded RNA-seq data is shown. **d**. Example of a genomic region with two convergent partially overlapping genes *PDCD4* and *BBIP1*, where transcripts from both strands are present in the oocyte cytoplasm but not in the oocyte nucleus. The cytoplasm contains spliced mRNA for both genes, whereas the nucleoplasm contains partially processed nascent RNA for only one gene. The exon-intron structure shown corresponds to one of the annotated transcript variants.

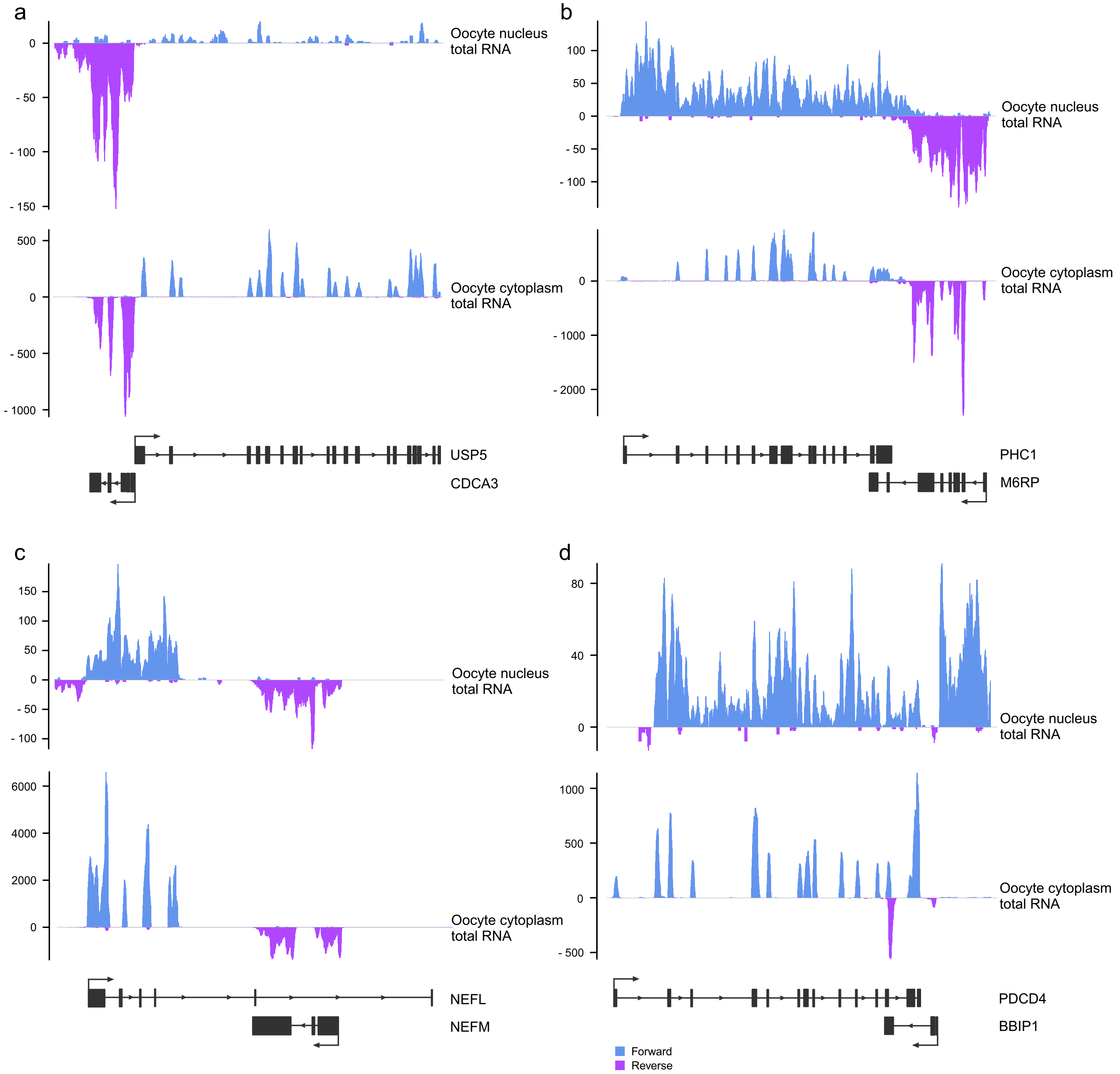
a
Oocyte nucleus total RNA
Oocyte cytoplasm total RNA
USP5
CDCA3
b
Oocyte nucleus total RNA
Oocyte cytoplasm total RNA
PHC1
M6RP
c
Oocyte nucleus total RNA
Oocyte cytoplasm total RNA
NEFL
NEFM
d
Oocyte nucleus total RNA
Oocyte cytoplasm total RNA
PDCD4
BBIP1
Forward
Reverse

**Figure S6. Correlation between poly(A) RNA and total RNA reads from (a) oocyte nucleus and (b) oocyte cytoplasm.**

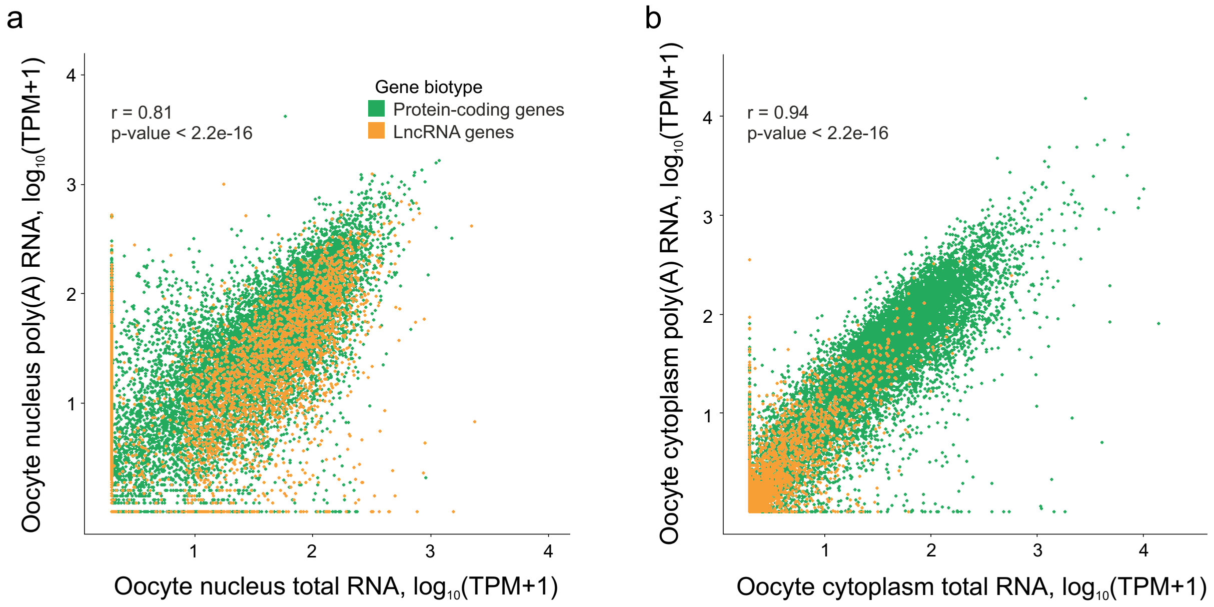
a
r = 0.81
p-value < 2.2e-16
Gene biotype
Protein-coding genes
LncRNA genes
Oocyte nucleus poly(A) RNA, log10(TPM+1)
Oocyte nucleus total RNA, log10(TPM+1)
b
r = 0.94
p-value < 2.2e-16
Oocyte cytoplasm poly(A) RNA, log10(TPM+1)
Oocyte cytoplasm total RNA, log10(TPM+1)

**Figure S7. Visualisation of nascent pre-mRNA from individual transcribed protein-coding genes (*CSMD3*, *INPPA5* and *PARD3B*) by RNA-FISH on lampbrush chromosomes.**

**a, b, c** - overview of the chromosomal regions of *CSMD3* (**a**), *INPPA5* (**b**) and *PARD3B* (**c**) genes with the positions of BAC clones used as RNA-FISH probes; **a', b', c'** - coverage tracks of total and poly(A) RNA from the oocyte nuclei (cyan) and cytoplasm (orange) of lampbrush stage oocytes visualised with IGV browser, data are shown for the short fragments covered by BAC clones, count range limits are given within square brackets. **a'', b'', c''** - RNA-FISH and with BAC clones CH261-97C18 (green) to *CSMD3** (**a''**), CH261-179F2 (yellow) to *INPPA5* (**b''**), and CH261-93F1 (red) to *PARD3B* (**c''**) genes on chicken lampbrush chromosomes 2 (**a''**), 6 (**b''**), and 7 (**c''**). Schematic drawings summarising several RNA-FISH micrographs indicate direction of transcription. Scale bars – 10 µm. DAPI – grayscale.

* Presumably, an unannotated transcript variant, approximately 77.1 kbp in length, is synthesised from the CSMD3 gene in lampbrush stage oocytes; this transcription unit makes up a short lateral loop with an average contour length of 3.3 µm.

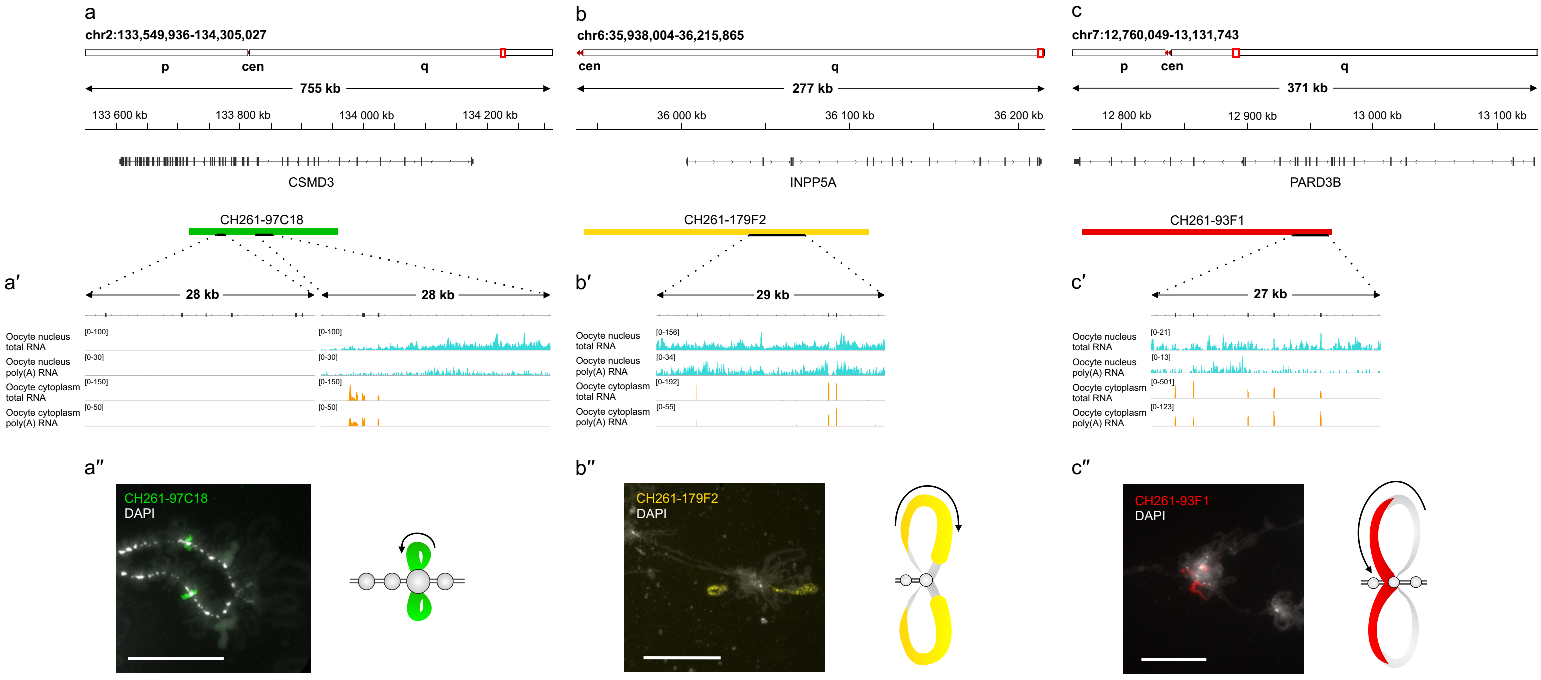
a
chr2:133,549,936-134,305,027
p
cen
q
755 kb
133 600 kb
133 800 kb
134 000 kb
134 200 kb
CSMD3
CH261-97C18
a′
28 kb
28 kb
Oocyte nucleus total RNA
Oocyte nucleus poly(A) RNA
Oocyte cytoplasm total RNA
Oocyte cytoplasm poly(A) RNA
[0-100]
[0-30]
[0-150]
[0-50]
a″
CH261-97C18
DAPI
b
chr6:35,938,004-36,215,865
cen
q
277 kb
36 000 kb
36 100 kb
36 200 kb
INPP5A
CH261-179F2
b′
29 kb
[0-156]
[0-34]
[0-192]
[0-55]
b″
CH261-179F2
DAPI
c
chr7:12,760,049-13,131,743
p
cen
q
371 kb
12 800 kb
12 900 kb
13 000 kb
13 100 kb
PARD3B
CH261-93F1
c′
27 kb
[0-21]
[0-13]
[0-501]
[0-123]
c″
CH261-93F1
DAPI

**Figure S8. At the post-lampbrush chromosome stage, the transcription loops with nascent gene transcripts become substantially shorter. a, b** - overview of the genomic regions containing transcribed genes *RBMS3* (**a**) and *CCSER1* (**b**) with the positions of BAC clones used as FISH probes. Coverage tracks of total RNA from the nuclei of lampbrush chromosome stage (cyan) and post-lampbrush chromosome stage (light cyan) oocytes visualised with IGV browser; count range limits are given within square brackets. **c-f** - visualisation of nascent pre-mRNA from individual transcribed protein-coding genes. RNA-FISH with BAC clone-based probes CH261-163C1 (green) and CH261-63A12 (red) to the beginning and to the end of the *RBMS3* gene on chicken chromosome 2 at lampbrush (**c**) and post-lampbrush stage (**d**). RNA-FISH with BAC clone-based probe CH261-124F10 (green) to the beginning of *CCSER1* gene on chicken chromosome 4 at lampbrush (**e**) and post-lampbrush stage (**f**). Chromosomes at lampbrush and post-lampbrush stages are shown at the same magnification. Labelled lateral loops shorten substantially at the post-lampbrush chromosome stage (arrows). Scale bars – 10 µm. DAPI – grayscale or blue.

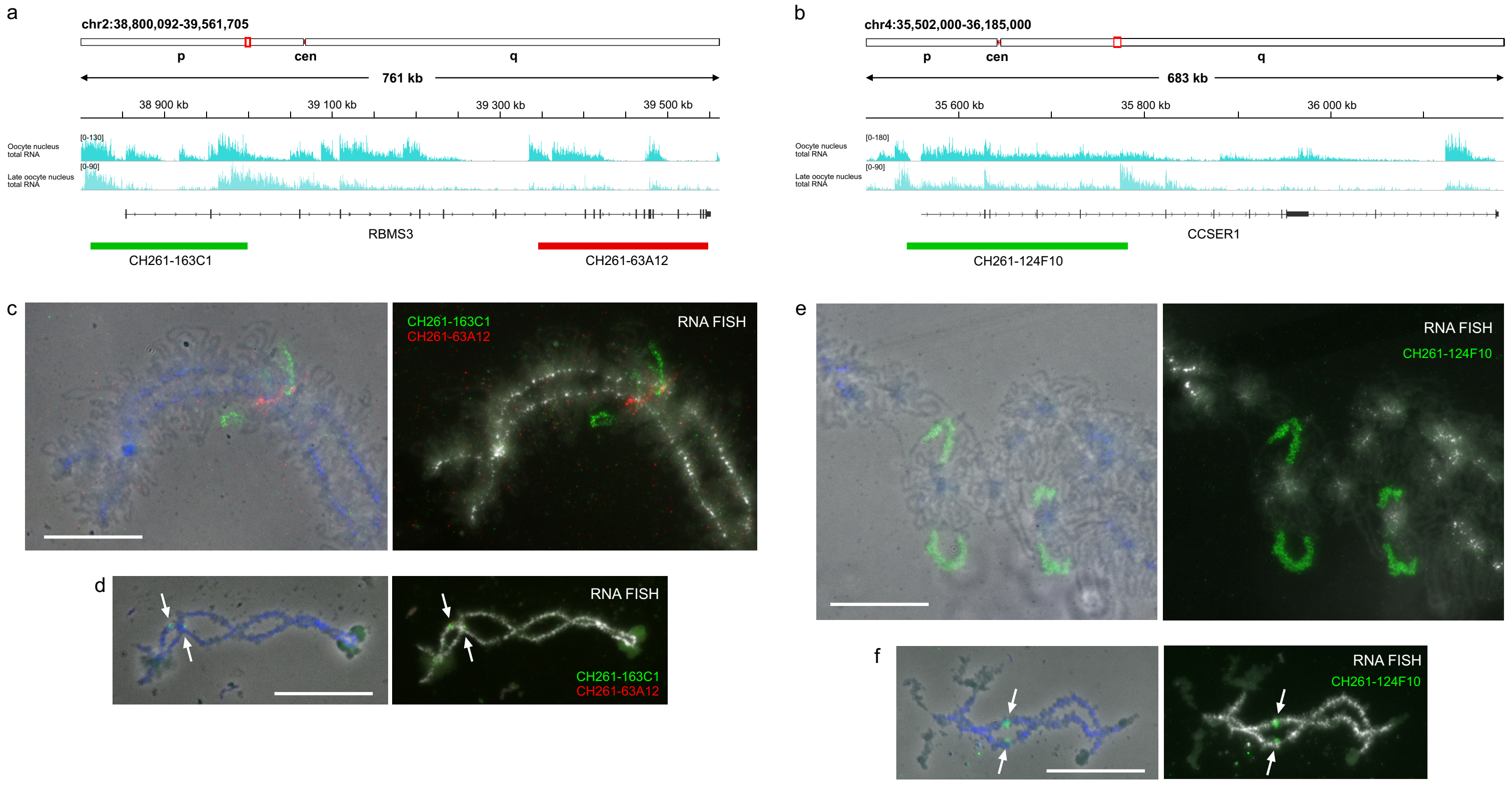
a
chr2:38,800,092-39,561,705
p
cen
q
761 kb
38 900 kb
39 100 kb
39 300 kb
39 500 kb
Oocyte nucleus total RNA
Late oocyte nucleus total RNA
RBMS3
CH261-163C1
CH261-63A12
b
chr4:35,502,000-36,185,000
p
cen
q
683 kb
35 600 kb
35 800 kb
36 000 kb
Oocyte nucleus total RNA
Late oocyte nucleus total RNA
CCSER1
CH261-124F10
c
CH261-163C1
CH261-63A12
RNA FISH
d
RNA FISH
CH261-163C1
CH261-63A12
e
RNA FISH
CH261-124F10
f
RNA FISH
CH261-124F10

**Figure S9. Comparison of RNA-seq data and RNA-FISH pattern along the genomic regions containing multiple transcribed genes and forming lateral loops.**

**a**-**c** - genomic regions on chicken chromosomes 1 (**a**), 6 (**b**) and 8 (**c**) covered by BAC clones used as RNA-FISH probes (bottom row); coverage tracks of total and poly(A) RNA from the oocyte nuclei (cyan) and cytoplasm (orange) of lampbrush stage oocytes visualised with IGV browser. Count range limits are given within square brackets. **a'**-**b'** - RNA-FISH with BAC clones CH261-104F19 (red) and CH261-54H10 (green) on chicken lampbrush chromosome 1 (**a'**), CH261-94G14 (green) on chicken lampbrush chromosome 6 (**b'**) and CH261-96D24 (red) on chicken lampbrush chromosome 8 (**c'**). Schematic drawings summarising several RNA-FISH micrographs are shown. Scale bars – 10 µm. DAPI – grayscale.

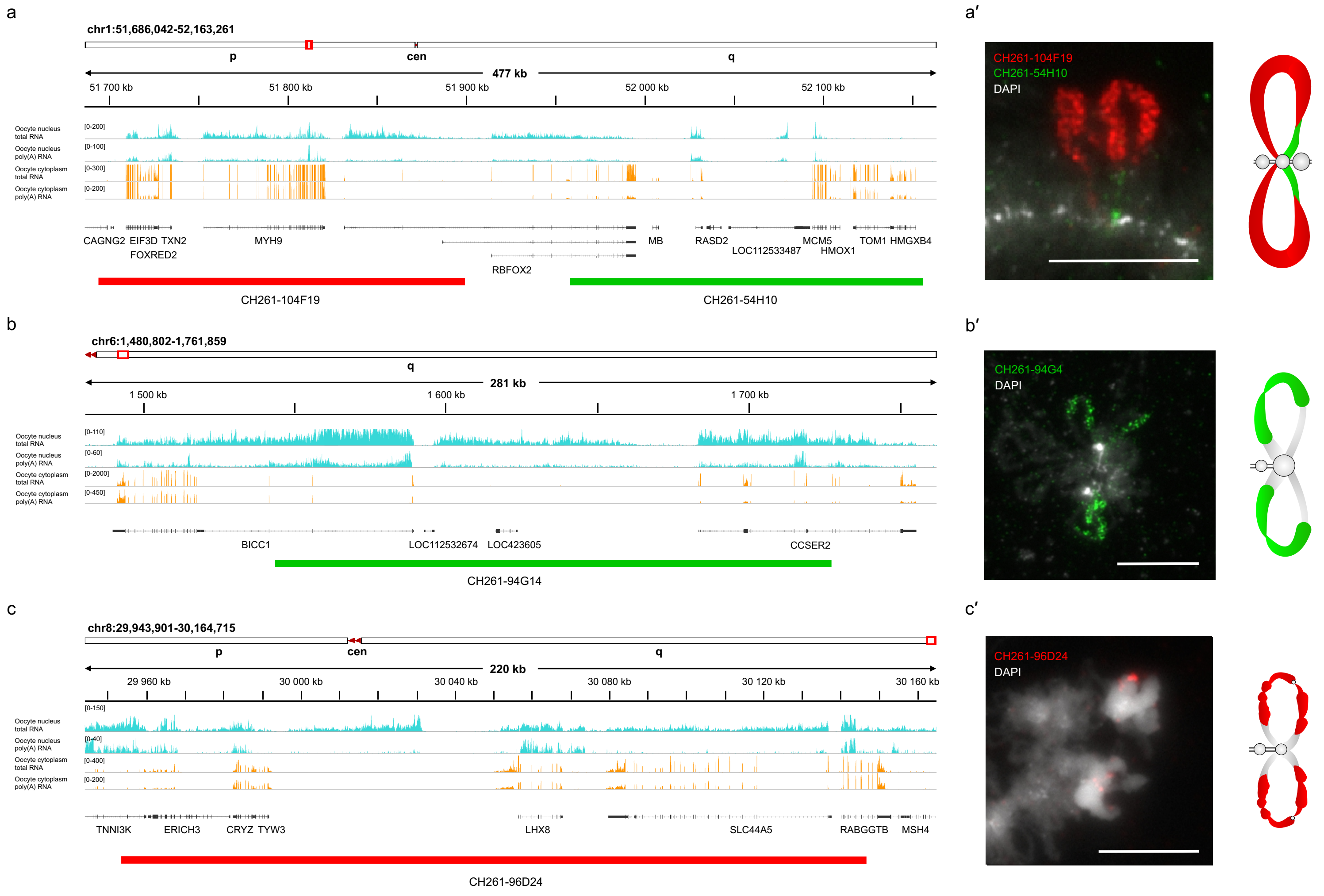
a
chr1:51,686,042-52,163,261
p
cen
q
477 kb
51 700 kb
51 800 kb
51 900 kb
52 000 kb
52 100 kb
Oocyte nucleus total RNA
Oocyte nucleus poly(A) RNA
Oocyte cytoplasm total RNA
Oocyte cytoplasm poly(A) RNA
CAGNG2 EIF3D TXN2
FOXRED2
MYH9
RBFOX2
MB
RASD2
LOC112533487
MCM5
HMOX1
TOM1 HMGXB4
CH261-104F19
CH261-54H10
a′
CH261-104F19
CH261-54H10
DAPI
b
chr6:1,480,802-1,761,859
q
281 kb
1 500 kb
1 600 kb
1 700 kb
BICC1
LOC112532674
LOC423605
CCSER2
CH261-94G14
b′
CH261-94G4
DAPI
c
chr8:29,943,901-30,164,715
p
cen
q
220 kb
29 960 kb
30 000 kb
30 040 kb
30 080 kb
30 120 kb
30 160 kb
TNNI3K
ERICH3
CRYZ TYW3
LHX8
SLC44A5
RABGGTB
MSH4
CH261-96D24
c′
CH261-96D24
DAPI

**Figure S10. Telomerase reverse transcriptase and telomerase RNA expression during oogenesis and early embryogenesis.**

**a, b.** Coverage tracks of RNA-seq data for lampbrush stage oocyte nucleus, post-lampbrush stage oocyte nucleus, oocyte cytoplasm, whole oocyte, zygote and embryonic stages EGK.I, III, VI, VIII, and X are shown for the telomerase reverse transcriptase (TERT) (**a**) and telomerase RNA (TERC) (**b**) genes, demonstrating preservation of the mature maternal telomerase RNAs from the oocyte to embryo stage EGK.VIII with the appearance of nascent zygotic transcripts at the EGK.X stage.

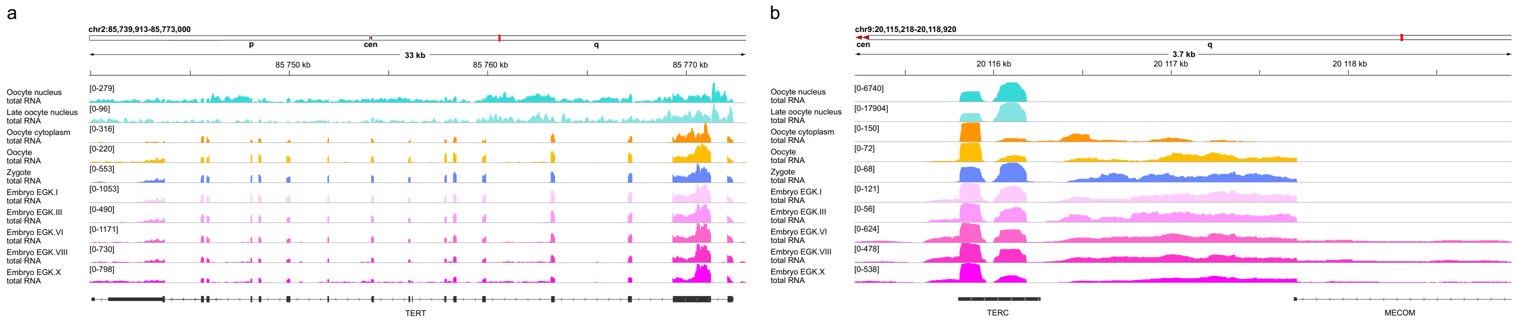

a
chr2:85,739,913-85,773,000
p
cen
q
33 kb
85 750 kb
85 760 kb
85 770 kb
Oocyte nucleus total RNA
[0-279]
Late oocyte nucleus total RNA
[0-96]
Oocyte cytoplasm total RNA
[0-316]
Oocyte total RNA
[0-220]
Zygote total RNA
[0-553]
Embryo EGK.I total RNA
[0-1053]
Embryo EGK.III total RNA
[0-490]
Embryo EGK.VI total RNA
[0-1171]
Embryo EGK.VIII total RNA
[0-730]
Embryo EGK.X total RNA
[0-798]
TERT
b
chr9:20,115,218-20,118,920
cen
q
3.7 kb
20 116 kb
20 117 kb
20 118 kb
Oocyte nucleus total RNA
[0-6740]
Late oocyte nucleus total RNA
[0-17904]
Oocyte cytoplasm total RNA
[0-150]
Oocyte total RNA
[0-72]
Zygote total RNA
[0-68]
Embryo EGK.I total RNA
[0-121]
Embryo EGK.III total RNA
[0-56]
Embryo EGK.VI total RNA
[0-624]
Embryo EGK.VIII total RNA
[0-478]
Embryo EGK.X total RNA
[0-538]
TERC
MECOM

**Figure S11. Examples of transcribing and non-transcribing small housekeeping RNA genes. a-c.** Two transcribing U6 snRNA genes (**a**), transcribing 5S rRNA gene (**b**) and non-transcribing U7 snRNA gene (**c**) are shown. **d-e.** Transcribing 7SK (**d**) and 7SL (**e**) RNA genes are shown. **f-g.** Examples of transcribed small housekeeping RNA genes with corresponding stable intronic sequence RNAs (sisRNAs) resistant to RNase R treatment. *HSPA8* host gene for embedded SNORD14 (**f**) and *NCAPD2* host gene for embedded U85 scaRNA (**g**) are shown. Coverage tracks of total RNA, poly(A) RNA and small RNA from the oocyte nuclei (cyan) and cytoplasm (orange) of lampbrush stage oocytes visualised by IGV browser along the gene body are shown; count range limits are given within square brackets. Coverage track of total RNA from the oocyte nuclei after RNase R treatment reveals sisRNAs (green). The exon-intron structure shown corresponds to one of the annotated transcript variants.

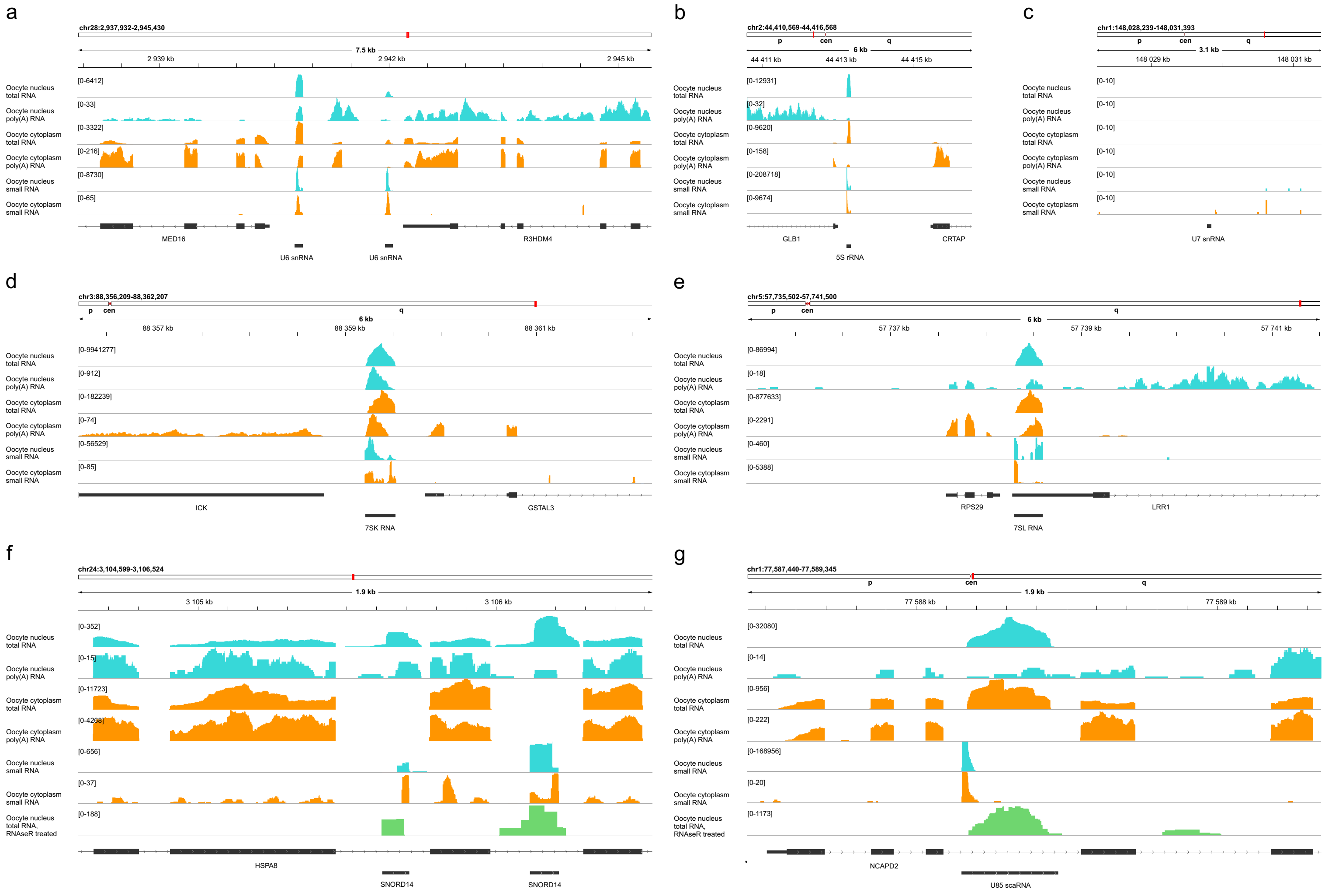
a
chr28:2,937,932-2,945,430
7.5 kb
2 939 kb
2 942 kb
2 945 kb
Oocyte nucleus total RNA
Oocyte nucleus poly(A) RNA
Oocyte cytoplasm total RNA
Oocyte cytoplasm poly(A) RNA
Oocyte nucleus small RNA
Oocyte cytoplasm small RNA
[0-6412]
[0-33]
[0-3322]
[0-216]
[0-8730]
[0-65]
MED16
R3HDM4
U6 snRNA
U6 snRNA
b
chr2:44,410,569-44,416,568
p
cen
q
6 kb
44 411 kb
44 413 kb
44 415 kb
[0-12931]
[0-32]
[0-9620]
[0-158]
[0-208718]
[0-9674]
GLB1
CRTAP
5S rRNA
c
chr1:148,028,239-148,031,393
3.1 kb
148 029 kb
148 031 kb
[0-10]
U7 snRNA
d
chr3:88,356,209-88,362,207
6 kb
88 357 kb
88 359 kb
88 361 kb
[0-9941277]
[0-912]
[0-182239]
[0-74]
[0-56529]
[0-85]
ICK
GSTAL3
7SK RNA
e
chr5:57,735,502-57,741,500
6 kb
57 737 kb
57 739 kb
57 741 kb
[0-86994]
[0-18]
[0-877633]
[0-2291]
[0-460]
[0-5388]
RPS29
LRR1
7SL RNA
f
chr24:3,104,599-3,106,524
1.9 kb
3 105 kb
3 106 kb
Oocyte nucleus total RNA, RNAseR treated
[0-352]
[0-15]
[0-11723]
[0-4268]
[0-656]
[0-37]
[0-188]
HSPA8
SNORD14
SNORD14
g
chr1:77,587,440-77,589,345
1.9 kb
77 588 kb
77 589 kb
[0-32080]
[0-14]
[0-956]
[0-222]
[0-168956]
[0-20]
[0-1173]
NCAPD2
U85 scaRNA

**Figure S12. Correlation between miRNA (a, b) and piRNA (c, d) reads in biological replicates from oocyte nucleus (a, c) and cytoplasm (b, d).**

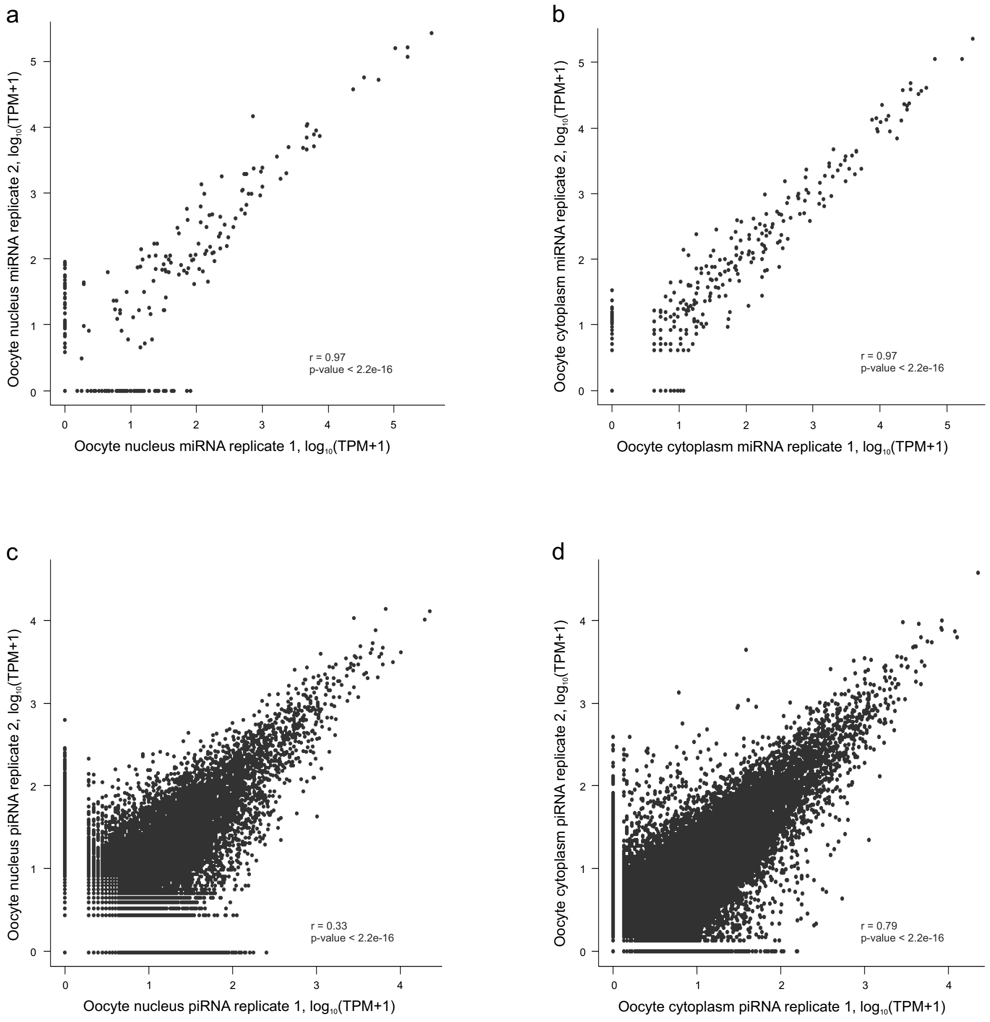

a
Oocyte nucleus miRNA replicate 2, log10(TPM+1)
Oocyte nucleus miRNA replicate 1, log10(TPM+1)
r = 0.97
p-value < 2.2e-16
b
Oocyte cytoplasm miRNA replicate 2, log10(TPM+1)
Oocyte cytoplasm miRNA replicate 1, log10(TPM+1)
r = 0.97
p-value < 2.2e-16
c
Oocyte nucleus piRNA replicate 2, log10(TPM+1)
Oocyte nucleus piRNA replicate 1, log10(TPM+1)
r = 0.33
p-value < 2.2e-16
d
Oocyte cytoplasm piRNA replicate 2, log10(TPM+1)
Oocyte cytoplasm piRNA replicate 1, log10(TPM+1)
r = 0.79
p-value < 2.2e-16

**Table S1.** RNA-seq data available from the NCBI Sequence Read Archive (SRA) using the accession numbers provided.

| **Sample** | **RNA library** | **Sample Name** | **SRA accession number** |
|---|---|---|---|
| Diplotene lampbrush stage oocytes from adult egg-laying females. Single oocyte cytoplasm was collected after nucleus isolation. | Poly(A) RNA library | Oop-1_polyA | SRR23800671 |
| Diplotene lampbrush stage oocytes from adult egg-laying females. Single oocyte cytoplasm was collected after nucleus isolation. | Poly(A) RNA library | Oop-2_polyA | SRR23800670 |
| Diplotene lampbrush stage oocytes from adult egg-laying females. Single oocyte cytoplasm was collected after nucleus isolation. | Small RNA library | Oop-1_small | SRR23800667 |
| Diplotene lampbrush stage oocytes from adult egg-laying females. Single oocyte cytoplasm was collected after nucleus isolation. | Small RNA library | Oop-2_small | SRR23800666 |
| Diplotene lampbrush stage oocytes from adult egg-laying females. Cytoplasm from 10 oocytes was collected after nucleus isolation. | Total RNA library | Oop_ctrl | SRR23800665 |
| Diplotene lampbrush stage oocytes from adult egg-laying females. 20 oocyte nuclei were isolated by micromanipulations. | Poly(A) RNA library | GV-K_polyA | SRR23800664 |
| Diplotene lampbrush stage oocytes from adult egg-laying females. 20 oocyte nuclei were isolated by micromanipulations. | Poly(A) RNA library | GV-M_polyA | SRR23800663 |
| Diplotene lampbrush stage oocytes from adult egg-laying females. 10 oocyte nuclei were isolated by micromanipulations. | Small RNA library | GV-1_small | SRR23800662 |
| Diplotene lampbrush stage oocytes from adult egg-laying females. 10 oocyte nuclei were isolated by micromanipulations. | Small RNA library | GV-2_small | SRR23800661 |
| Diplotene lampbrush stage oocytes from adult egg-laying females. 20 oocyte nuclei were isolated by micromanipulations. | Total RNA library | GV1-LBC_RNA | SRR23800660 |
| Diplotene lampbrush stage oocytes from adult egg-laying females. 10 oocyte nuclei were isolated by micromanipulations. RNA was treated by RNase R. | Total RNA library | GV_RNAse_R | SRR23800669 |
| Diplotene post-lampbrush stage oocytes from adult egg-laying females. 10 oocyte nuclei were isolated by micromanipulations. | Total RNA library | GV1-large_RNA | SRR23800668 |

**Table S2.** The list of BAC clones containing fragments of chicken genomic DNA from the CHORI-261 library that were used as DNA-probes for FISH. Coordinates are indicated according to galGal6 chicken genome assembly; green highlighting indicates BAC-clones that overlap with the sequence of only one annotated gene.

| **Chromosome region (Mb)** | **BAC clone name** | **Start coordinate (bp)** | **End coordinate (bp)** | **Insert length (bp)** | **Labelled with** | **Figure # with the FISH-mapping data** |
|---|---|---|---|---|---|---|
| GGA1_50-52 | CH261-104F19 | 51694570 | 51899748 | 205179 | dig | Figure S9 a′ |
| GGA1_50-52 | CH261-54H10 | 51958408 | 52156056 | 197649 | bio | Figure S9 a′ |
| GGA1_148-149 | CH261-191J12 | 148978858 | 149168480 | 189623 | bio | Figure 4 c′′, c′′′ |
| GGA1_148-149 | CH261-90G18 | 149755715 | 149967613 | 211899 | dig | Figure 4 c′′, c′′′ |
| GGA2_38-40 | CH261-163C1 | 38813307 | 39000127 | 186821 | dig | Figure 4 a′′<br>Figure S8 c, d |
| GGA2_38-40 | CH261-63A12 | 39346898 | 39548608 | 201711 | bio | Figure 4 a′′<br>Figure S8 c, d |
| GGA2_133-134 | CH261-97C18 | 133718475 | 133959233 | 240759 | bio | Figure S7 a′′ |
| GGA2_143 | CH261-17J16 | 143308662 | 143489517 | 180856 | bio | Figure 4 d′′, d′′′ |
| GGA4_35-37 | CH261-124F10 | 35545929 | 35782990 | 237062 | bio | Figure 4 b′′<br>Figure S8 e, f |
| GGA4_35-37 | CH261-109C8 | 36773434 | 37003277 | 229844 | dig | Figure 4 b′′ |
| GGA6_1 | CH261-94G14 | 1543421 | 1727121 | 183701 | bio | Figure S9 b′ |
| GGA6_35-36 | CH261-179F2 | 35942801 | 36112050 | 169250 | Atto-647 | Figure S7 b′′ |
| GGA7_12-13 | CH261-93F1 | 12768085 | 12968061 | 199977 | dig | Figure S7 c′′<br>Figure 6 b, c |
| GGA7_12-13 | CH261-126G14 | 13073010 | 13332073 | 259064 | bio | Figure 6 b, c |
| GGA7_12-13 | CH261-38J23 | 13383341 | 13562115 | 178775 | dig | Figure 6 b, c |
| GGA8_29-30 | CH261-96D24 | 29953369 | 30146659 | 193291 | dig | Figure S9 c′ |
| GGA14_11-12 | CH261-179I1 | 11242090 | 11428383 | 186294 | dig | Figure 5 b-b′ |
| GGA14_11-12 | CH261-119E18 | 11515602 | 11713554 | 197953 | bio | Figure 5 b-b′ |
| GGA14_11-12 | CH261-78O7 | 11849039 | 12022809 | 173771 | Atto-647 | Figure 5 b-b′ |

**Table S3.** The list of annotated gene sequences (according to galGal6 chicken genome assembly) whose transcriptional status was verified by both RNA-FISH on lampbrush chromosomes and RNA-seq of oocyte nuclear and cytoplasmic RNA.

| **Gene name** | **Chromosome location** | **BAC clone name** | **Transcriptional status** | **Figure # with the FISH-mapping data** |
|---|---|---|---|---|
| *RBMS3* | chr2:38,853,312-39,552,113 (+) | CH261-163C1, CH261-63A12 | Transcribed | Figure 4 a-a′′ |
| *CSMD3* | chr2:133606128-134178021 (-) | CH261-97C18 | Transcribed | Figure S7 a-a′′ |
| *CCSER1* | chr4:35560875-36180105 (+) | CH261-124F10 | Transcribed | Figure 4 b-b′′ Figure S8 e |
| *GRID2* | chr4:36337862-37032793 (+) | CH261-109C8 | Transcribed | Figure 4 b-b′′ |
| *PARD3B* | chr7:12762049-13129743 (-) | CH261-93F1 | Transcribed | Figure 6 a-b Figure S7 c-c′′ |
| *RBFOX1* | chr14:11257243-12010152 (-) | CH261-179I1, CH261-119E18, CH261-78O7 | Transcribed | Figure 5 a-b′′ |
| *GPC5* | chr1:148802059-149423920 (-) | CH261-191J12 | Non-transcribed | Figure 4 c-c′′′ |
| *LOC112531542 (ncRNA)* | chr1:149,914,589-149,919,702 (-) | CH261-90G18 | Non-transcribed | Figure 4 c-c′′′ |
| *THSD7A* | chr2:26258257-26528121 (-) | CH261-96F4 | Transcribed | Figure 3 in Kulikova *et al.* 2022 |
| *DIAPH2* | chr4:5797966-5955595 (-) | CH261-177B7 | Transcribed | Figure 5 in Kulikova *et al.* 2022 |
| *COL25A1* | chr4:37348440-37647585 (+) | CH261-49F17 | Transcribed | Figure 26.1 in Kulikova and Krasikova 2022 |
| *KIFC3* | chr11:481558-491211 (+) | WAG52K20 | Transcribed | Figure 4 a in Zlotina *et al.* 2012 |

**Table S4.** Targets of predicted piRNAs in the chicken oocyte nucleus and oocyte cytoplasm.

| Repeat class | Repeat family | Oocyte nucleus replicate 1, % of reads | Oocyte nucleus replicate 2, % of reads | Oocyte cytoplasm replicate 1, % of reads | Oocyte cytoplasm replicate 2, % of reads | Oocyte nucleus replicate 1, unique piRNAs | Oocyte nucleus replicate 2, unique piRNAs | Oocyte cytoplasm replicate 1, unique piRNAs | Oocyte cytoplasm replicate 2, unique piRNAs |
|---|---|---|---|---|---|---|---|---|---|
| DNA transposon | DNA/Crypton-A | 0,000 | 0,000 | 0,000 | 0,000 | 0 | 0 | 0 | 0 |
| | DNA/Kolobok | 0,000 | 0,000 | 0,000 | 0,000 | 0 | 0 | 0 | 0 |
| | DNA/PIF-Harbinger | 0,000 | 0,000 | 0,000 | 0,000 | 0 | 0 | 0 | 0 |
| | DNA/TcMar-Mariner | 0,142 | 0,151 | 0,042 | 0,105 | 210 | 137 | 489 | 1111 |
| | DNA/hAT-Blackjack | 0,000 | 0,000 | 0,000 | 0,000 | 0 | 0 | 0 | 0 |
| | DNA/hAT-Charlie | 0,015 | 0,037 | 0,016 | 0,057 | 21 | 31 | 127 | 326 |
| | DNA/Unknown | 0,027 | 0,029 | 0,006 | 0,016 | 38 | 28 | 77 | 170 |
| LINE | LINE/CR1 | 40,069 | 43,964 | 54,512 | 49,821 | 22053 | 16100 | 79557 | 88497 |
| | LINE/Penelope | 0,000 | 0,000 | 0,000 | 0,000 | 0 | 0 | 0 | 1 |
| | LINE/R2 | 0,008 | 0,018 | 0,002 | 0,001 | 10 | 18 | 38 | 12 |
| LTR | LTR | 0,002 | 0,000 | 0,000 | 0,000 | 2 | 0 | 0 | 0 |
| | LTR/ERV | 0,137 | 0,124 | 0,103 | 0,114 | 186 | 106 | 659 | 801 |
| | LTR/ERV1 | 0,099 | 0,078 | 0,088 | 0,040 | 128 | 74 | 677 | 351 |
| | LTR/ERVK | 0,910 | 1,011 | 0,657 | 1,617 | 889 | 672 | 2880 | 5445 |
| | LTR/ERVL | 18,664 | 16,759 | 29,408 | 29,286 | 8736 | 5565 | 37526 | 36581 |
| | LTR/Ngaro | 0,000 | 0,000 | 0,000 | 0,000 | 0 | 0 | 0 | 0 |
| | LTR/Pao | 0,000 | 0,000 | 0,000 | 0,000 | 0 | 0 | 0 | 0 |
| | LTR/Unknown | 3,481 | 3,779 | 2,435 | 3,130 | 2160 | 1484 | 5628 | 7501 |
| SINE | SINE/5S-Deu-L2 | 0,000 | 0,000 | 0,000 | 0,000 | 0 | 0 | 0 | 0 |
| | SINE/MIR | 0,007 | 0,000 | 0,000 | 0,000 | 9 | 0 | 0 | 2 |
| | SINE/tRNA | 0,000 | 0,000 | 0,000 | 0,000 | 0 | 0 | 0 | 0 |
| tRNA | tRNA | 0,000 | 0,000 | 0,000 | 0,000 | 0 | 0 | 6 | 3 |
| Unclassified repeats | | 5,755 | 7,051 | 7,693 | 9,082 | 4012 | 3334 | 13801 | 18266 |
| Reads aligning to a non-repetitive region | | 30,684 | 26,998 | 5,038 | 6,729 | 29885 | 14537 | 16391 | 31643 |

**List of References for the Supplementary Materials**

**List of Supplementary Data Sets**

**Supplementary Data Set 1.** RNA-seq data of levels of gene transcripts in total RNA samples from chicken oocyte nucleus and oocyte cytoplasm.
